# Multi-modal and multi-region distance model for neuroimaging: Application to ABCD study

**DOI:** 10.64898/2026.01.21.700689

**Authors:** Xinyu Zhang, Simon Vandekar, Andrew A. Chen, Kaidi Kang, Jakob Seidlitz, Aaron Alexander-Bloch, Jinyuan Liu

## Abstract

Large-scale neuroimaging studies often collect multiple modalities, such as task and resting-state functional MRI, diffusion MRI, and structural MRI. Joint inference across these modalities uses shared variation to improve statistical efficiency, increase replicability, and provide a more integrated view of brain-phenotype associations. In practice, however, such analyses are limited because complex cross-modality covariance cannot be flexibly modeled, which makes the resulting joint effects difficult to interpret. A recent distance-based ANOVA extension allows multimodal analysis and increases power for detecting group differences, but it cannot easily distinguish location from scale effects in distance space, offers only an omnibus pseudo-*F* test without interpretable parameters, and requires computationally intensive permutation inference. We propose a novel semiparametric, U-statistics-based Generalized Estimating Equation (UGEE) framework that unifies univariate and multivariate distance models. By regressing pairwise dissimilarities on covariates, this method yields interpretable regression coefficients that disentangle location and scale effects and quantify inter-modality differences, while flexibly accounting for correlations among modality distances. The estimator is based on efficient influence functions, ensuring asymptotic efficiency, robustness to misspecification, and computational scalability for large-scale data analysis. We evaluate the proposed method through extensive simulations and analyses of the Adolescent Brain Cognitive Development dataset. Results show that UGEE accurately estimates modality, group, and interaction effects and achieves a 100-fold speed-up compared with permutation-based approaches. This framework provides a general and computationally efficient tool for semiparametric inference on multimodal data, particularly suited for large neuroimaging applications.

## 1. Introduction

Understanding how brain features differ across diagnostic or developmental subgroups is a central goal of neuroimaging research (Van Rooij et al., 2018; Laurent et al., 2020; Lin, Si and Kang, 2024; Ginestet et al., 2017). However, the high dimensionality, complex dependence, and non-Gaussian nature of imaging data, such as voxel-wise structural features or functional connectivity matrices, pose substantial challenges for conventional multivariate analysis of variance (MANOVA) approaches (Krzanowski, 2006). In response, distance-based methods such as the Permutational Multivariate Analysis of Variance (PERMANOVA) (Anderson, 2001; Anderson and Braak, 2003), or equivalently, Multivariate Distance Matrix Regression (MDMR), have emerged as powerful tools for detecting group-level differences in complex, high-dimensional outcomes (Reiss et al., 2010; Alexander-Bloch et al., 2012). These methods directly focus on the dissimilarity space, partitioning total variation in pairwise distances into between- and within-group components and yielding a pseudo-*F* statistic as a global measure of association (Liu et al., 2024). By aggregating information across thousands of imaging features into a single distance matrix, they bypass the multiple comparisons that plague mass-univariate analyses and have therefore become a powerful and versatile tool for detecting group-level differences in both structural MRI features and functional connectivity profiles derived from resting-state fMRI data (Shinohara et al., 2020).

Recent neuroimaging initiatives have increasingly collected multiple imaging modalities from the same participants, enabling the joint analysis of complementary structural and functional information. For instance, the Adolescent Brain Cognitive Development (ABCD) Study is a large-scale longitudinal project that follows 11,868 youth across 21 U.S. sites, beginning at ages 9-10 and continuing through adolescence (Marek et al., 2022; Tomasi and Volkow, 2024). It includes both structural and functional MRI, among other assessments, which provide unprecedented opportunities to study multimodal neural development. The rise of these datasets has driven a surge of methodological advances in multimodal integration (Gao et al., 2020). One recent approach, similarity-based multimodal regression (SiMMR), extends MDMR to model multiple imaging modalities simultaneously through their pairwise distance profiles (Chen et al., 2024). By leveraging information across modalities, SiMMR improves the power to detect group-level differences compared with single-modality analyses and has become a promising direction for nonparametric multimodal inference.

Despite their success, existing distance-based approaches have several limitations. First, PERMANOVA and MDMR are inherently nonparametric ANOVA-type tests (Anderson and Braak, 2003); they provide a single pseudo-*F* statistic and an omnibus p-value without yielding interpretable regression coefficients or predictive mappings between predictors and outcomes. Consequently, these frameworks do not easily quantify the direction or magnitude of effects, which may hinder the scientific interpretability and preclude their use in power analysis for a new study design. Second, these methods often conflate location and scale effects in the distance space (Anderson, 2001; Liu et al., 2022a), making it difficult to distinguish mean differences from differences in dispersion or variability (Warton, Wright and Wang, 2012). Third, the permutation-based inference underlying these methods can be computationally prohibitive for large-scale datasets such as ABCD (e.g., SiMMR takes 10.6 minutes to run on *n* = 854 in our data application). Finally, while SiMMR increases sensitivity by summing distances across modalities, this aggregation may overlook dependence among repeated modality measures and obscure the ability to detect modality-specific or cross-modality interaction effects (Lahat, Adali and Jutten, 2015). Determining which modalities carry the most informative signal to diagnostic or developmental distinctions is not only scientifically meaningful, but also practically important, given the high cost of multimodal imaging acquisition (approximately US$1,000 per hour; Marek et al. (2022)).

In this paper, we develop a semiparametric U-statistics-based Generalized Estimating Equation (UGEE) framework for inference that unifies multivariate imaging features under a distance-based regression paradigm. The proposed model directly regresses pairwise dissimilarities on covariates across subjects, analogous to how the conventional GEE characterizes the regression relationships. Hence, our model yields interpretable regression coefficients and effect sizes that can be used for inference, prediction, and study design. By incorporating efficient influence functions (EIF), the estimator achieves asymptotic efficiency (Tsiatis, 2006; Kennedy, 2018), robustness to distributional misspecification, and computational scalability even in very large samples. We further extend this framework to parameterize multiple modalities and allow flexible modeling of both modality-specific and cross-modality interaction effects while accounting for correlations among modalities. Together, these advances bridge the gap between nonparametric distance-based testing and semiparametric regression modeling.

The rest of the paper is organized as follows. Section 2 introduces the distance-based analysis framework for both single- and multi-modality. We start by reviewing the standard distance-based ANOVA and distance-based regression in the single-modality setting. Then, for the multi-modality setting, we present a novel unified distance-based regression method in contrast with the existing ANOVA-type approaches. We present comprehensive simulation studies in Section 3 and apply the proposed model to the ABCD dataset to investigate associations between multiple imaging modalities and key phenotypic measures in Section 4. We give concluding remarks in Section 5.

## 2. Methods

### 2.1. Single Modality

We start by reviewing standard distance approaches for a single modality. Consider a sample of *n* subjects and let **y**_*i*_ denote a *p ×* 1 column vector of multivariate outcomes for the *i*th subject (*i* = 1, 2, · · ·, *n*). In most neuroimaging studies, the feature size *p* can be large, which may exceed the sample size. For example, **y**_*i*_ can represent structural phenotypes such as the cortical thickness or gray matter volume, where *p*, referred to as “feature size,” represents the number of voxels, brain regions, or parcellation units.

Motivated by comparing group differences in the multivariate neuroimaging outcome **y**_*i*_, we begin by constructing pairwise distances between subjects. For any pair of subjects 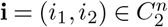, we define pairwise distance 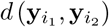 based on a chosen distance or dissimilarity metric, where 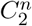 denotes the set of all unordered pair combinations from the integer set {1, …, *n*}. This strategy integrates information across all features within this modality without explicit feature selection, thereby mitigating potential information loss (Liu et al., 2023). A wide range of metrics *d*(·, ·) may be used, including Euclidean, Jaccard, and other dissimilarity measures (Hancock, 2004). These upper-triangular entries uniquely determine the symmetric pairwise distance matrix (*d*^*y*^)_*n×n*_, with the lower-triangular entries obtained by symmetry and zeros along the diagonal. This *n × n* matrix serves as the basis for the downstream analysis; a summary of all notations is provided in Supplement Section S5.1.

#### 2.1.1. Distance-based ANOVA

A straightforward inferential framework for the distance outcome *d*^*y*^ is the Permutational Multivariate Analysis of Variance (PERMANOVA), or the generalized Multivariate Distance Matrix Regression (MDMR) (Anderson, 2001; Reiss et al., 2010). This framework is particularly useful when *x*_*i*_ indexes (sub)groups (1 ≤ *x*_*i*_ ≤ *K*). Assuming no additional covariates, let *µ*_*k*_ and 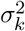 denote the mean and variance of pairwise outcome distance within the *k*th subgroup. The null hypothesis of equal group means can be expressed as:

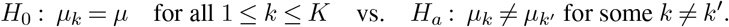

PERMANOVA performs ANOVA on distances by partitioning total dissimilarity into within- and between-group components. Let **A** = −0.5(*d*^*y*^) denote the scaled distance matrix, and define the Gower-centered matrix **G** = (**I** − **11**^⊤^*/n*)**A**(**I** − **11**^⊤^*/n*), where **I** and **1** denote the identity matrix and an *n* × 1 vector of ones, respectively.

Let **H** = **X**(**X**^⊤^**X**)^−1^**X**^⊤^ denote the projection (“hat”) matrix corresponding to the design matrix **X**, consisting of an intercept and group indicators (e.g., dummy variables). A pseudo-*F* statistic is then defined as

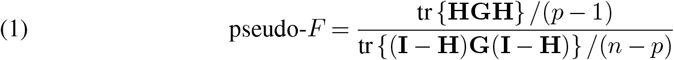

where tr{·} is the trace function. Since the distance is not restricted to Euclidean, the test statistic in (1) does not generally follow an exact *F* -distribution, and its significance is assessed via nonparametric permutation tests (Anderson and Braak, 2003). Following Reiss et al. (2010), the degrees of freedom can be omitted as they remain constant across permutations.

This framework readily extends to include continuous covariates in **X** and has been widely adopted in disciplines such as ecology, neuroimaging, and microbiome research (Anderson, 2001; Reiss et al., 2010). Although commonly referred to as a “regression” in the MDMR literature, this framework is conceptually closer to an ANOVA in dissimilarity space than to a conventional regression model. The key distinction is the absence of an explicit (semi)parametric conditional mean mapping between predictors *x*_*i*_ and outcomes *d*^*y*^ (Tsiatis, 2006). Instead, MDMR evaluates whether distances among multivariate responses differ systematically across levels or linear combinations of **X** by partitioning the centered distance (McArtor, Lubke and Bergeman, 2017).

Consequently, MDMR does not yield interpretable regression coefficients nor support prediction for new observations; the design matrix **X** serves only to encode group contrasts or covariate-induced partitions of the dissimilarity space. As such, MDMR provides a global omnibus test, analogous to an ANOVA *F* -test, without quantifying effect sizes or estimating mappings between predictors and outcomes. Moreover, MDMR can be computationally intensive and often exhibits limited power for detecting between-group difference, particularly “location” type of effects (Warton, Wright and Wang, 2012; McArtor, Lubke and Bergeman, 2017; Liu et al., 2022a; Shi et al., 2023). To address these limitations, we introduce a more flexible distance-based regression framework next.

#### 2.1.2. Distance-based Regression

In the same vein of subgroup comparison within a distance-based regression framework, we first map each subject-level covariate *x*_*i*_ to a between-subject attribute. For a given pair 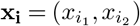, we construct *pairwise* indicators, or dummy variables, through a *one-hot encode* function:

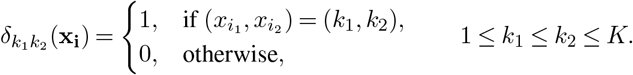

Here, 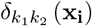 indicates pairs with *concordant* subgroup membership (*k*_1_ = *k*_2_ = *k*) or *discordant* membership (*k*_1_ *< k*_2_). Collecting all such indicators yields

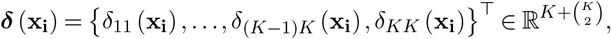

which enumerates all possible concordant and discordant subgroup combinations, with 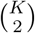 denoting the number of discordant pairs that can be formed from *K* distinct levels.

For example, if *x*_*i*_ represents diagnosis (Disease vs. Healthy), then 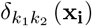 corresponds to three levels {*δ*_DD_ (**x**_**i**_), *δ*_HH_ (**x**_**i**_), *δ*_HD_ (**x**_**i**_)}, indexing Disease-Disease, Healthy-Healthy, and mixed pairs. We designate one as the reference (e.g., *δ*_DD_ (**x**_**i**_)) and add other levels to the linear predictor. Then, the distance regression comparing subgroups becomes

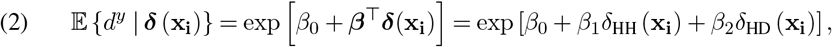

with exp(*β*_1_) representing the mean difference in *d*^*y*^ comparing Healthy-Healthy vs. Disease-Disease pairs and exp(*β*_2_) that between mixed vs. Disease-Disease pairs.

Compared with the ANOVA-based framework, this regression enables granular characterization of heterogeneous patterns in *d*^*y*^ (Liu et al., 2022a). It naturally accommodates multiple covariates of mixed types, including high-dimensional features or covariates available only through pairwise distances or kernels, and provides a flexible mechanism for integrating such information without explicit low-dimensional embeddings (Liu et al., 2024). Moreover, the regression setup induces a natural basis for modeling repeated measures and within-subject dependence, which we discuss next.

### 2.2. Multi-modality

To enhance the characterization of brain organization, multiple neuroimaging modalities are increasingly collected for the same participants within a single study (Biessmann et al., 2011). Denote by **y**_*im*_ a *p*_*m*_ *×* 1 vector of imaging outcomes for *i*th subject from the *m*th modality (*m* = 1, …, *M*). For example, the structural modality may include regional cortical thickness or gray matter volume, whereas the functional modality may include vectorized functional connectivity values (Kaczkurkin, Raznahan and Satterthwaite, 2019; Tagliazucchi and Laufs, 2015). In both cases, the feature dimension may greatly exceed the sample size (*p*_*m*_ ≫ *n*).

Beyond high dimensionality, these repeated imaging modalities exhibit strong within-cluster correlation, analogous to the dependence modeled in univariate repeated-measures ANOVA, mixed-effects models, or generalized estimating equations (GEE). Therefore, valid inference requires considerations on both dimensionality and within-cluster dependence (Liang and Zeger, 1986).

To address the high dimensionality inherent to each modality, we again transform imaging features using (dis)similarity metrics (Krzanowski, 2006). Specifically, let 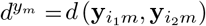 denote the pairwise dissimilarity metric for the *m*th modality, evaluated for subject pair 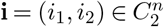. Collecting these quantities yields the vector

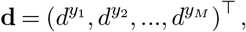

which represents the multi-modal dissimilarity profile for the **i**th subject pair across all *M* modalities.

#### 2.2.1. Distance-based ANOVA

Within the ANOVA framework, a recent line of work on similarity-based multimodal regression (SiMMR) extended classical PERMANOVA in (1) from a single to multiple modalities by combining modality-specific distance profiles (Chen et al., 2024). For each modality *m*, the pairwise distance matrix is first centered to obtain **G**_*m*_ = (**I** − **11**^⊤^*/n*)**A**_*m*_(**I** − **11**^⊤^*/n*), where 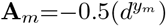 and 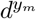 denotes the modality-specific dissimilarity matrix. The overall similarity structure is then formed by summing across modalities, yielding 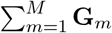.

A pseudo-*F* statistic is obtained by substituting this aggregated matrix for **G** in (1). Under the null hypothesis, observations are exchangeable, and inference proceeds via permutation of the hat matrix **H** = **X**(**X**^⊤^**X**)^−1^**X**^⊤^, with degrees of freedom omitted (Chen et al., 2024).

Further, investigators may be interested in assessing the contribution of a subset of covariates **X**_*r*_ ⊂ **X**. To this end, a Dempster’s trace-type statistic (Dempster, 1958), referred to as SiMMR-D, has been proposed:

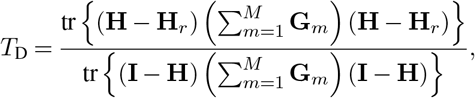

where 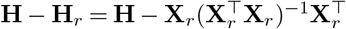 corresponds to the reduced model excluding **X**_*r*_.

While SiMMR-D accommodates multimodal distance information, it implicitly assumes independence across modalities by aggregating (dis)similarity matrices via summation. To account for inter-modality correlation, classical multidimensional scaling (cMDS) is first applied within each modality to obtain low-dimensional coordinate representations **Z**_*m*_ (Anderson, 2001), such that 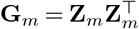. These modality-specific scores are then concatenated across modalities, and principal component analysis (PCA) is applied to stabilize rank and reduce dimensionality. The resulting PC scores therefore capture shared variation and correlations across modalities. Associations between the covariate subset **X**_*r*_ and the multimodal imaging outcomes are subsequently tested using the SiMMR-PC statistic based on Pillai’s trace:

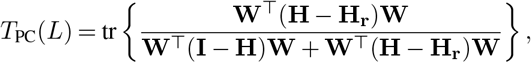

where **W** ∈ R^*n×L*^ contains the first *L* PC scores.

Although these SiMMR-based approaches extend distance-based ANOVA to the multimodal setting and can accommodate repeated distance measures, they inherit key limitations of permutation-based inference. In particular, they do not yield interpretable effectsize parameters, impose substantial computational burden, and may exhibit limited power, as demonstrated in our simulation studies. Motivated by these limitations, we develop in the next section two semiparametric distance-based regression frameworks that directly estimate effect sizes and support fully asymptotic inference without permutation.

#### 2.2.2. Distance-based Regression

Regressions naturally incorporate covariates with different types. Let *x*_*i*_ denote (sub)group membership for the subject *i*, and suppose that *q* additional covariates are available. To preserve their original data structures, we partition these covariates into continuous and categorical components, (**z**_*i*_, **w**_*i*_), where 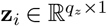 contains continuous covariates and 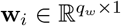 contains categorical covariates, with *q* = *q*_*z*_ + *q*_*w*_. Throughout, **w**_*i*_ refers to additional categorical covariates other than the group indicator *x*_*i*_.

For continuous covariates, we construct pairwise distances 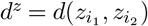 across all subject pairs, yielding a 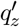 -dimensional vector 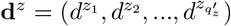, where 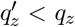. This allows dimension reduction when appropriate; for example, multiple continuous covariates may be grouped to form composite distance measures.

Categorical covariates **w**_*i*_ are incorporated by constructing pairwise dummy variables ***δ***(**w**_**i**_), analogous to the group indicator ***δ***(**x**_**i**_) introduced in Section 2.1.2. Together, these constructions enable mixed-type covariates to be incorporated within a unified distance-based regression framework.

Extending the ANOVA framework to distance-based regression further enables flexible modeling of high-dimensional and multimodal outcomes, with distance responses modeled either univariately or multivariately, depending on the application. We start with the univariate model.

##### 2.2.2.1. Univariate Distance-Based Regression (UGEE-Uni)

Analogous to SiMMR-D, the average of all modality-specific distances can be treated as a univariate outcome. The conditional expectation can be specified through the regression model

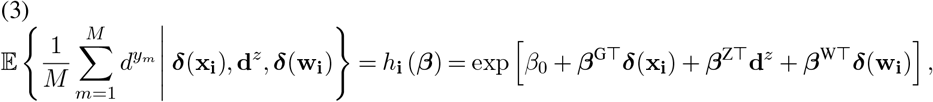

where ***β***^G^ captures the main group effects of interest, ***β***^Z^ and ***β***^W^ represent the effects from continuous and categorical covariates, respectively.

For example, exp(***β***^Z^) represents the multiplicative change in the expected average distance across the *M* modalities associated with a one-unit increase in **d**^*z*^, whereas exp(***β***^W^) measures the multiplicative effect attributable to differences defined by categorical covariates.

As such, the UGEE-Uni model provides a parsimonious summary of joint multimodal signal and is well suited for inference targeting global associations across modalities. However, by collapsing modality-specific distances through averaging, it does not capture intermodality dependence and cannot assess modality-specific effects or cross-modality interactions. These limitations can be addressed by model multimodal distances as a multivariate outcome, which we develop next.

##### 2.2.2.2. Multivariate Distance-Based Regression (UGEE-Multi)

We introduce a repeateddistance regression that jointly models all *M* modalities. Modality-specific effects are explicitly parameterized through indicator functions, yielding the following marginal mean model for each modality *m* = 1, …, *M* :

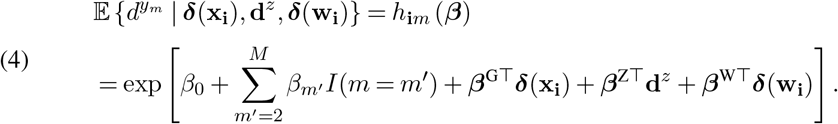

In the linear predictor, the first modality (*m* = 1) serves as the reference level with mean exp(*β*_0_), and *β*_*m*_*′* captures modality-specific mean differences relative to this reference. Thus, exp(*β*_*m*_*′*) can be interpreted as the ratio of the expected distance for modality *m*^*′*^ to that of the reference modality, holding all covariates fixed.

The coefficient vectors ***β***^G^, ***β***^Z^, and ***β***^W^ represent the main (sub)group effects and the effects of continuous and categorical covariates, respectively. As in UGEE-Uni, all covariate effects can be interpreted multiplicatively on the expected distance scale.

When the modality-by-group (**x**_**i**_) interactions are of interest, the model can be extended by adding

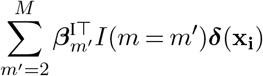

to the linear predictor of (4), where ***β***^I^ captures modality-specific heterogeneity in subgroup effects. Higher-order interactions with additional covariates can be incorporated analogously.

### 2.3. Semiparametric Efficient Inference

Classical pseudo-*F* statistics in the multivariate ANOVA framework do not, in general, follow a limiting *F* distribution (Shinohara et al., 2020). As a result, inference typically relies on computationally intensive permutation procedures, which can suffer from substantial power loss under certain alternatives (Liu et al., 2022b,a). Despite some prior attempts to derive its asymptotic distributions using spectral decompositions (McArtor, Lubke and Bergeman, 2017; Shi et al., 2023), these approaches impose restrictive conditions, such as requiring the distance matrix to be positive semi-definite, and even then, the analytical p-values still require approximation and may be conservative (McArtor, Lubke and Bergeman, 2017). Consequently, such methods have seen limited practical adoption; permutation-based inference remains the default in most R packages.

We address these limitations by developing a semiparametric inference framework grounded in the efficient influence function (EIF). This approach enables valid, fully asymptotic inference while improving efficiency and power and retaining substantial modeling flexibility (Kennedy, 2022).

A fundamental challenge in inference for distance outcomes arises from the dependence structure induced by pairwise construction. Specifically, pairwise distances 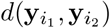 are not mutually independent: pairs that share a common subject are correlated. For example, *d*(**y**_1_, **y**_2_) and *d*(**y**_1_, **y**_3_) are correlated through **y**_1_. Independence holds only for pairs with disjoint subject indices, yielding a sparse but nontrivial correlation structure across all pairs (Liu et al., 2022b).

Let each subject’s observed data be denoted by 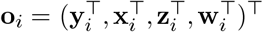, drawn independently from a population distribution *p* (**O**_*i*_; ***θ***_0_). The parameter vector is ***θ*** = (***β***^⊤^, *η*)^⊤^, where ***β*** is the finite-dimensional parameter of interest (e.g., as in (3) or (4)) and *η* represents an infinite-dimensional nuisance parameter, assumed variationally independent of ***β*** following standard semiparametric theory (Tsiatis, 2006). Semiparametric models allow the nuisance *η* to be unspecified, thereby enhancing robustness and flexibility in estimation of ***β***. Our goal is to derive efficient estimators and valid variance expressions for ***β*** ∈ ℝ^*q*^ using information from all subject pairs. To this end, the multivariate distance-based regression model across *M* modalities (UGEE-Multi) can be written as

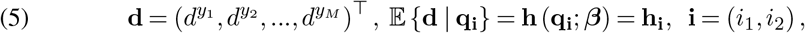

where **h**_**i**_ = (*h*_**i**1_, *h*_**i**2_, …, *h*_**i***M*_)^⊤^ denotes the vector of modality-specific mean functions and **q**_**i**_ = {***δ***(**x**_**i**_), **d**^*z*^, ***δ***(**w**_**i**_)} collects all transformed pairwise predictors.

Model (5) encompasses several special cases. When *M* = 1, it reduces to the single-modality model in (2); when 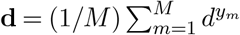, it corresponds to the UGEE-Uni model in (3). Throughout, the (joint) distribution of **d** is left unspecified and treated as an infinite-dimensional nuisance.

#### 2.3.1. Semiparametric Efficient Influence Function (EIF)

We introduce an influence-function-based semiparametric estimator which, despite involving 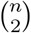 distinct pairs in the estimation of the target parameter ***β*** in (5), remains 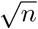 -consistent. The main result is summarized below. We include a sketch of the proof in the Supplemental Section S5.2.

##### Theorem 1.

*Let the true parameters be* 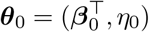, *under mild regularity conditions, the semiparametric efficient estimator* 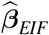 *based on all pairwise observations is* 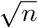 *consistent, asymptotic linear, and follows asymptotic normal (AN) distribution:*

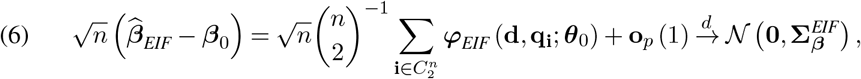

*with the efficient influence function (EIF)*

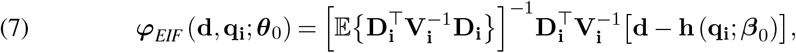

*where*

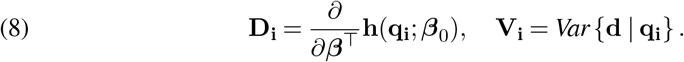

*This EIF leads to the smallest asymptotic variance within the model class defined by* (5):

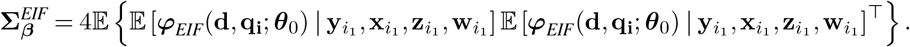

#### 2.3.2. Efficient UGEE

To implement the EIF-based estimator, we adopt a class of efficient *U-statistics-based Generalized Estimating Equations* (UGEE) for the pairwise outcomes. Because the variance-covariance structure **V**_**i**_ = Var (**d** | **q**_**i**_) in (8) is typically unknown, we can specify a working variance (Liang and Zeger, 1986):

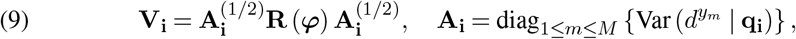

with **R** (***φ***)^*M×M*^ the working correlation matrix to account for potential dependence among the *m* modalities in 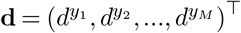.

For each subject pair 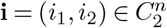, the UGEE is defined as

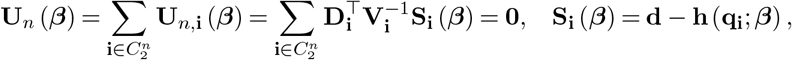

which uses the EIF in (7) and hence, yields an efficient estimator whose variance admits a “sandwich” form:

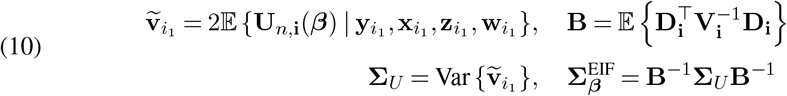

A consistent estimator of 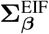 can be obtained by substituting consistent estimators of ***β*** and the corresponding moment estimators in (10). We outline the algorithm to obtain 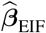 and 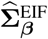 numerically as follows.

##### Algorithm 1: UGEE using EIF

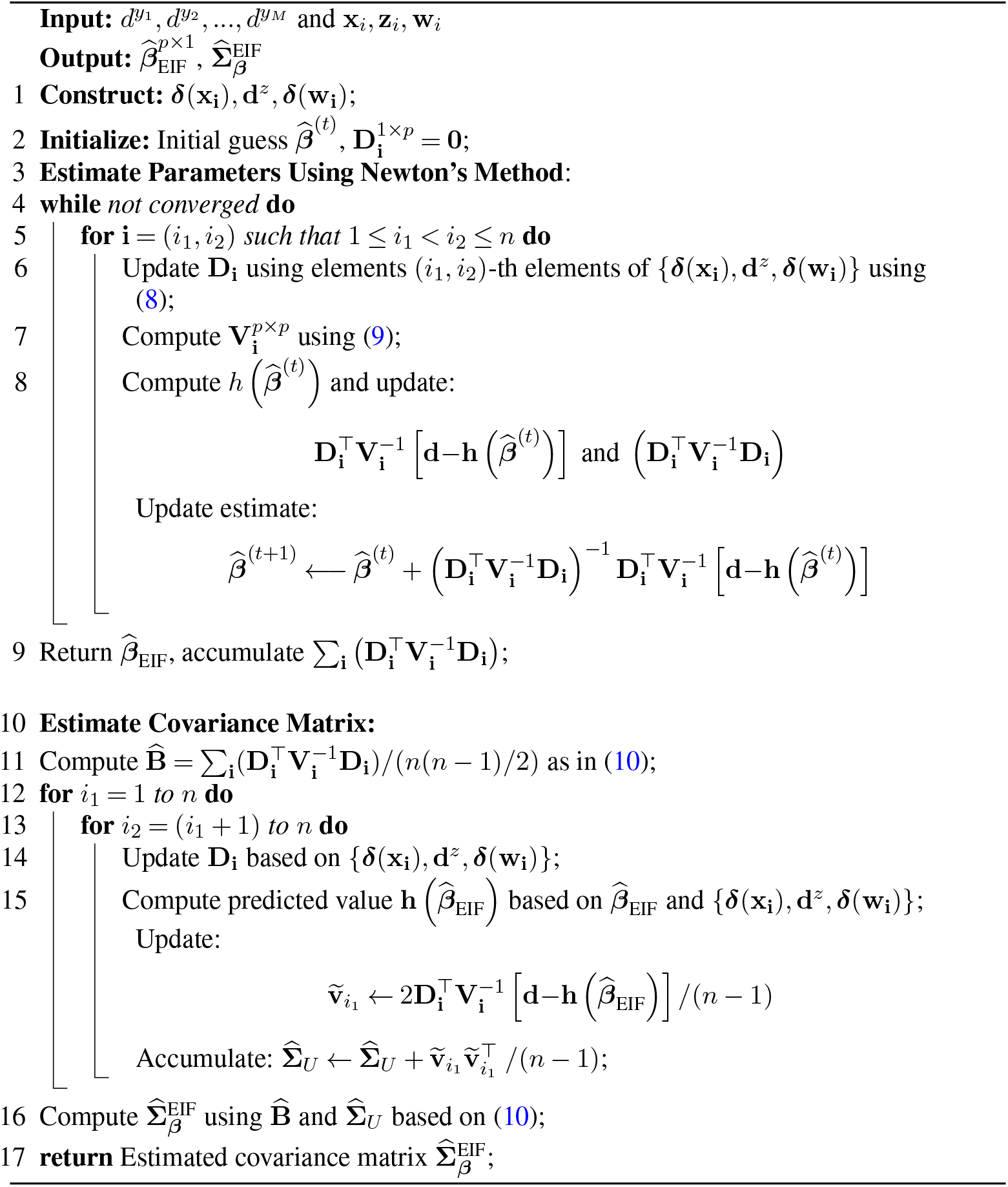

#### 2.3.3. Wald Test

For any hypotheses regarding the coefficients in (3) or (4), inference can be conducted using a Wald test based on the asymptotic distribution of the EIF-based estimator 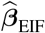 in (6). Specifically, the null hypothesis is first expressed through a linear contrast matrix **C** of rank *c*. The corresponding Wald statistic is then defined as

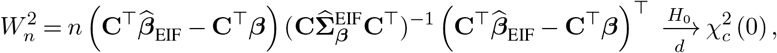

the limiting distribution of *W* ^2^ under the null *H*_0_ : **C**^⊤^***β*** = 0 is a central *χ*^2^ distribution with *c* degrees of freedom. This Wald test enables fully asymptotic inference without reliance on permutation and, by leveraging the efficient influence function, achieves high power for detecting true effects, as demonstrated in our simulation studies.

## 3. Simulation

We now evaluate the finite-sample performance of the proposed UGEE framework for detecting subgroups and modality effects, in comparison with existing alternatives such as SiMMR (using both Dempster and PC statistics).

### 3.1. Data-generating Mechanism

We considered two subgroups with *x*_*i*_ ∈ {1, 2} generated from Bernoulli(0.5). Then for each subject pair, we defined within-group and between-group indicators,

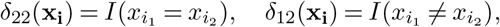

representing scale-type and location-type group differences, respectively.

For each subject, we generated three modalities (*m* = 1, 2, 3), each containing *p*_*m*_ imaging features. To induce correlations among features within and across modalities, we specified a 3 *×* 3 modality-level correlation structure. The within-modality correlation *ρ*_*w*_ was set to either 0 or 0.5, representing independent or moderately correlated features within each modality. The between-modality correlation *ρ*_*b*_ was set to 0, 0.2, or 0.8, corresponding to independent, weakly dependent, or strongly dependent exchangeable correlation structures across modalities.

Each block of the resulting correlation matrix was expanded to match the feature dimension of each modality, yielding

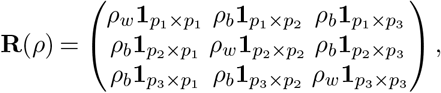

with diagonal elements fixed at 1. Conditions guaranteeing positive definiteness of **R**(*ρ*) are provided in Supplementary Section S5.3.

Next, residual errors in each modality, denoted by ***ϵ*** = (***ϵ***_1_, ***ϵ***_2_, ***ϵ***_3_), were drawn from a multivariate normal distribution MVN (**0**, *s*^2^**R**(*ρ*)), where *s* = 0.2 is the residual standard error. Next, we converted each residual matrix 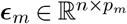 into a distance matrix 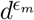 using Euclidean distance, and specify modality-by-group interactions

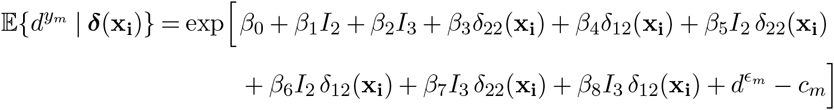

where *I*_2_ and *I*_3_ denote indicators for the second and third modalities. This yields the modality-specific distance outcomes 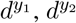, and 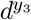.

Here the small correction *c*_*m*_ is used to offset the upward bias introduced due to the residual variance, which centers the baseline expectation at 1 under the null (Liu et al., 2022a). The closed-form expression of *c*_*m*_ is provided in the Supplement Section S5.4 under the assumption of feature independence (*ρ*_*w*_ = 0) and is empirically estimated with *n* = 5, 000 when features are correlated (*ρ*_*w*_ = 0.8).

### 3.2. Simulation Scenarios

We examined four types of data-generating scenarios comparing the SiMMR and UGEE (Uni- and Multi-):

(1) Null: No group or modality effects.
(2) Modality-only effects: Mean differences across modalities (second, or both second and third vs. reference).
(1) Group-only effects: Either scale (*δ*_22_) or location (*δ*_12_) effects between subgroups.
(4) Modality-and-group additive effects: Both modality and group effects contribute additively. Since SiMMR is not designed to capture modality-by-group interactions, we simulated an additional setting to compare between UGEE-Uni and UGEE-Multi to evaluate:
(5) Modality-by-group interaction effects: Group differences vary across modalities.

The true nonzero regression coefficients for each scenario are summarized in Table 1. In the first four scenarios, modalities were generated independently (*ρ*_*b*_ = 0), with within-modality correlation evaluated at two levels, *ρ*_*w*_ = 0 and 0.8, to assess performance under increasing feature dependence.

**Table 1.**
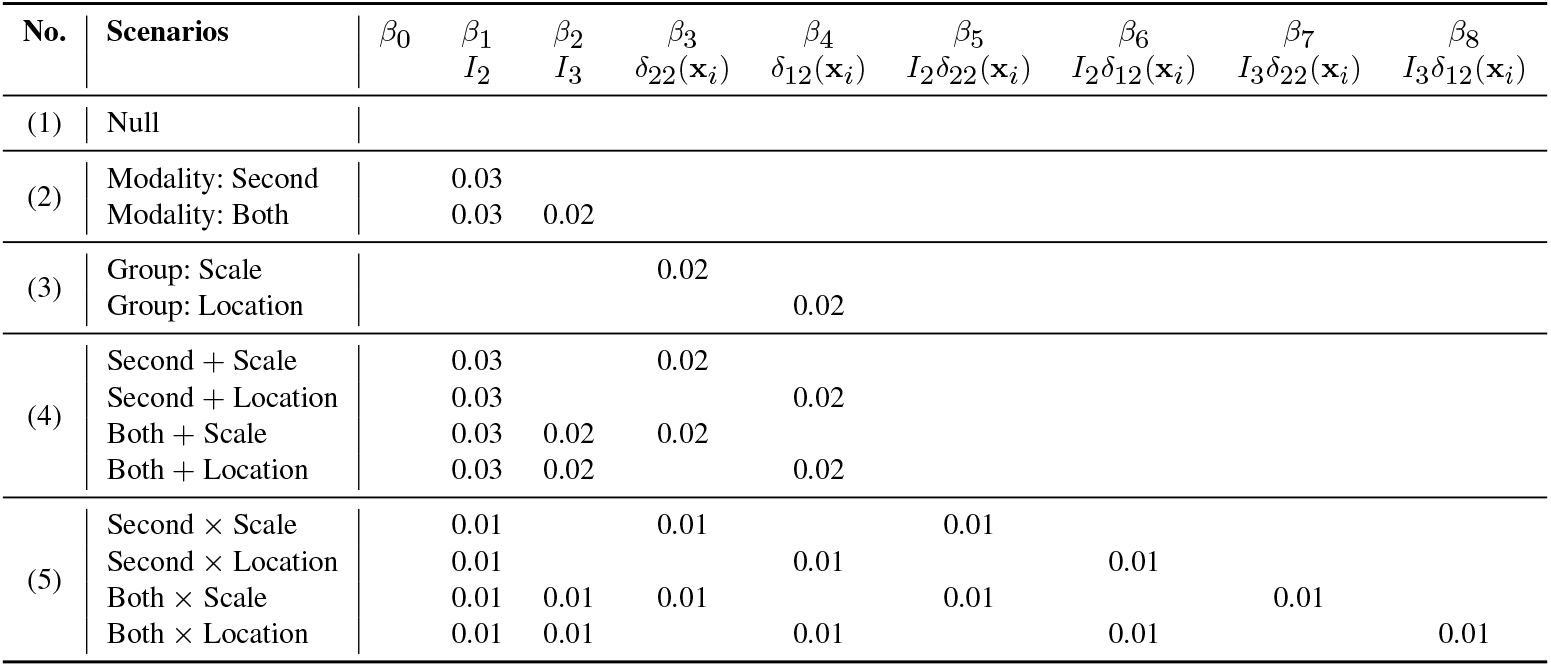
Specification of non-zero regression coefficients β under each simulation scenario. The second row of the header indicates the corresponding term for each coefficient. Coefficients with a value of zero are omitted from the table.

For computational comparability across methods, the number of permutations used in SiMMR was capped at 100. This choice reflects a pragmatic trade-off between computational feasibility and inferential resolution, consistent with common practice in large-scale simulation studies where permutation-based methods are known to incur substantial runtime (McArtor, Lubke and Bergeman, 2017). For SiMMR-PC, the PC corresponding to a positive eigenvalue was selected to maximize the total number of explained variability.

In the fifth scenario, we explicitly examined the robustness of UGEE to working-correlation misspecification. Data were generated with varying between-modality correlations (*ρ*_*b*_ = 0, 0.2, 0.8; *ρ*_*w*_ = 0.8) while UGEE models were fitted assuming both independence and exchangeable working correlation structures. For a sample size of 200, the resulting transformed distance correlations were approximately 0.65, 0.72, and 0.89, representing moderate to strong inter-modality dependence.

For each setting, we varied the sample size (*n* = 50, 75, 100, 150, 200) and performed 500 Monte Carlo replications. We evaluated: Type I error under the null (proportion of replications with *p* ≤ 0.05), statistical power under the alternatives, and bias of regression coefficients 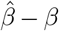, together with their standardized effect size 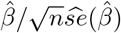. Both omnibus (global) and component-specific *a priori* hypothesis tests were assessed.

### 3.3. Simulation Results

As shown in Figure 1(A), all methods demonstrated satisfactory control of Type I error across tested hypotheses, although minor variability was observed at smaller sample sizes (*n* = 50). Due to aggregation across modalities, neither SiMMR nor UGEE-Uni detected modality differences, with power curves remaining near the nominal 0.05 line in Figure 1(B). This limitation arises because both models omit explicit parameterization of modality-specific effects. In contrast, UGEE-Multi consistently achieved higher power in scenarios where true modality effects exist, directly highlighting the benefit of modeling modalities as repeated distance outcomes rather than collapsing them into a single composite metric.

In general, both the permutation-based SiMMR and the semiparametric UGEE showed adequate performance in omnibus testing once the sample size exceeded moderate levels (e.g., *n >* 100). As expected, omnibus tests within UGEE achieved higher power than component-specific hypotheses (Figure 1(C) and (D)). Interestingly, SiMMR-PC showed instability in the presence of location effects: its power decreased with increasing sample size, e.g., from 94% when *n* = 50 to 84% when *n* = 100 and to 52% when *n* = 150 in Figure 1(C) location effect. This trend likely reflects limited information captured by a small number of retained PCs in small samples, and it persisted under additive modality effects (Figure 1(D)).

**FIG 1.**
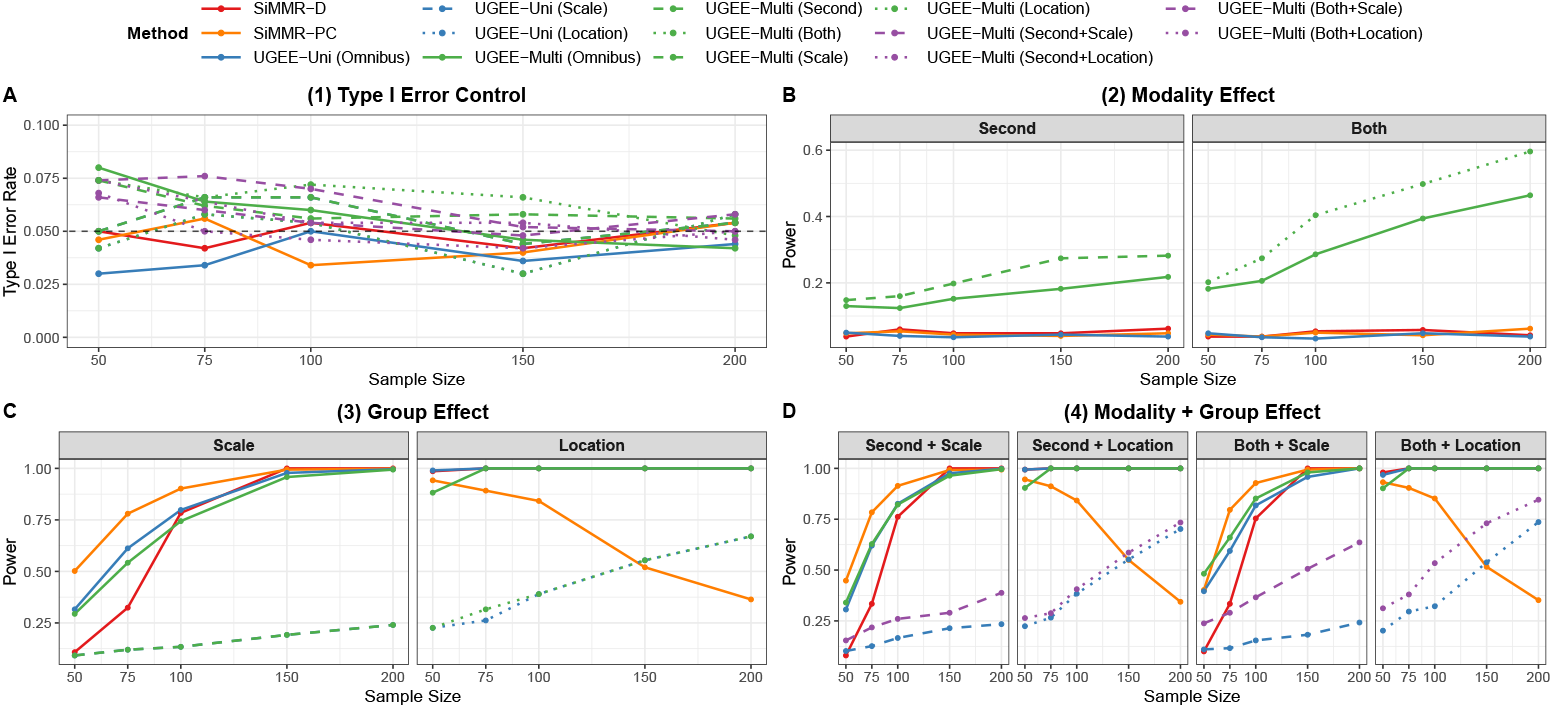
Power and type I error across different scenarios (sample size 50-200) with independent features within modality. (A) Type I error control under the null hypothesis. (B-D) Power for detecting modality differences, group differences, and combined effects, respectively. Colors represent method families (red: SiMMR-D, orange: SiMMR-PC, blue: UGEE-Uni, green: UGEE-Multi, purple: UGEE-Multi additive). Solid lines = omnibus tests (detect any difference); dashed/dotted lines = component-specific tests (designed for particular effects). Horizontal dashed line at 0.05 indicates the target Type I error rate.

When the within-modality correlation *ρ*_*w*_ was increased from 0 to 0.8, the effective information per modality decreased, inflating variance and slightly elevating the Type I error of omnibus tests for *n* ≤ 100 (see Figure 2(A)). Under such high correlation and moderate *n*, a nonparametric bootstrap may offer more reliable inference (Efron, 1979). Across all scenarios, UGEE-Uni exhibited declining power with increasing *ρ*_*w*_ (e.g., Figure 1(C) vs. 2(C)), reflecting its inability to offset information loss under stronger within-modality dependence. In contrast, UGEE-Multi benefited from a higher *ρ*_*w*_, e.g., comparing Figure 1(B) vs. 2(B), the omnibus power increased from 39% to 67% when *n* = 150, with both modalities specified to be different from the reference. This notable power gain is likely attributable to enhanced within-modality synchrony and amplified coherent signals across modalities, which allow the joint model to borrow strength across modalities. Coefficient estimates from both UGEE-Uni and UGEE-Multi remained consistent across settings; estimation accuracy summarized by coefficient bias and effect size is reported in Tables S2 and S3 in the Supplement.

**FIG 2.**
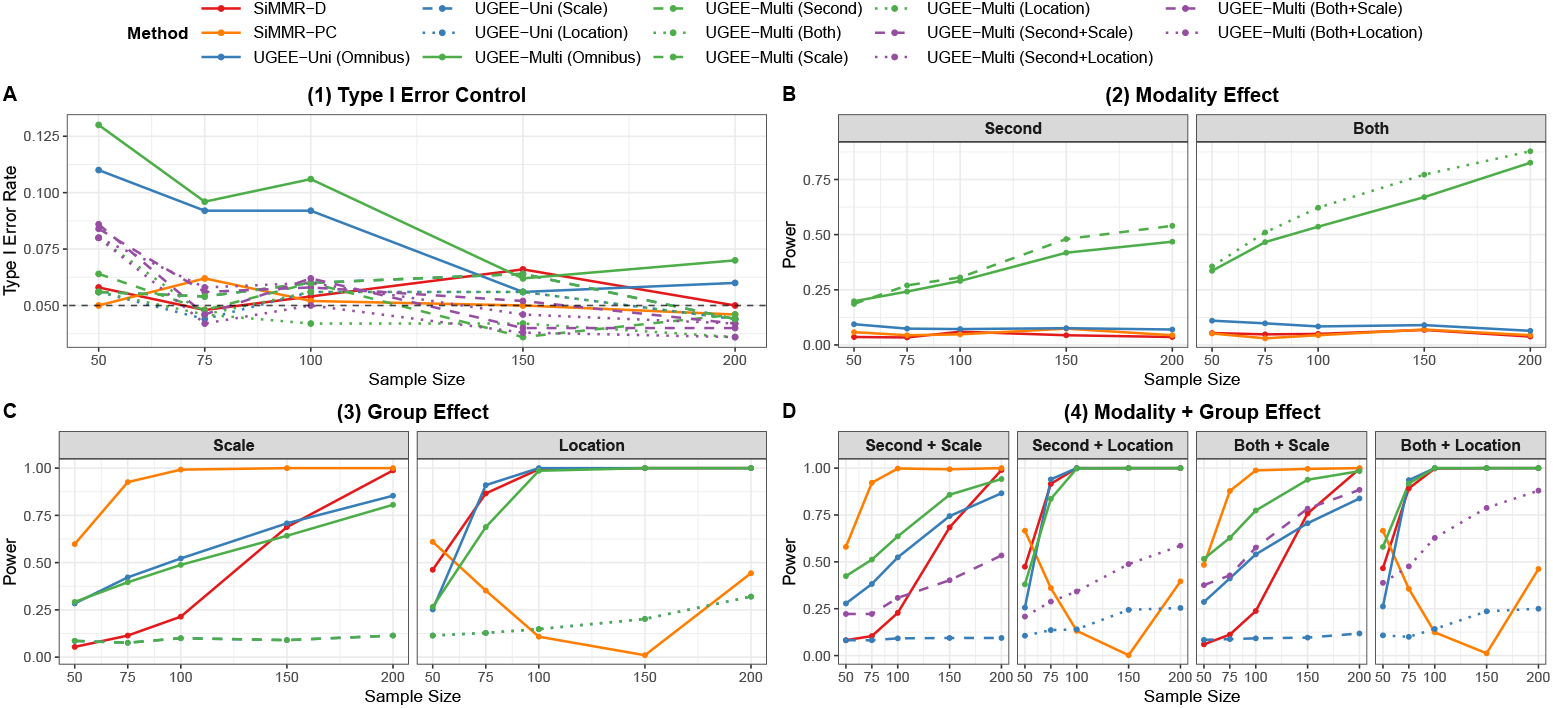
Power and type I error across different scenarios (sample size 50-200) with correlated features within modality. Same pattern as the above figure.

To further evaluate the impact of specifying working-correlation structures among modalities within the UGEE framework, Figure 3 presents power curves under data-generating processes with independent, moderately, and highly correlated inter-modality structures. UGEE-Multi showed strong robustness to working-correlation misspecification: power remained stable whether independence or exchangeable structures were assumed, thanks to the sandwich variance estimation. Moreover, the benefits of multimodal integration increased with stronger true inter-modality correlation, confirming that when modalities share covariance, joint modeling via UGEE-Multi can efficiently borrow strength to enhance statistical power. For example, with *n* = 50, power for detecting a location-by-both-modality effect increased from 65% to 76% as *ρ*_*b*_ increased from 0.2 to 0.8. Lastly, although omnibus tests capture multiple parameters and therefore yield higher overall power, component-specific tests remain valuable for targeted hypotheses, such as group-level scale or location effects.

**FIG 3.**
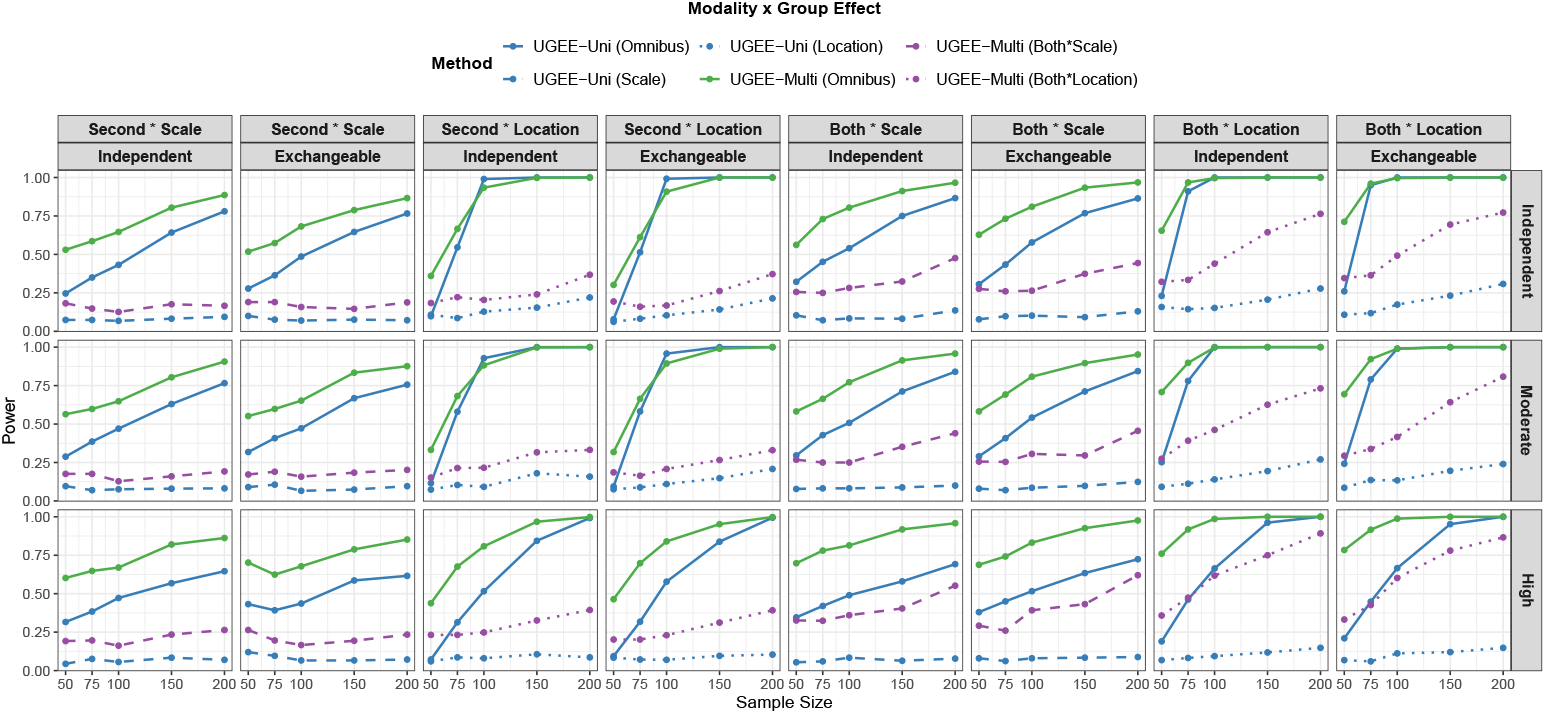
Power Across Different Testing Scenarios (UGEE only, sample size 50-200). Colors represent method families (blue: UGEE-Uni, green: UGEE-Multi, purple: UGEE-Multi interaction effects). Solid lines = omnibus tests (detect any difference); dashed/dotted lines = component-specific tests (designed for particular effects).

## 4. Application

### 4.1. Motivating ABCD Study

With nearly 12,000 participants aged 9-10 years at baseline, the ABCD Study provides an unparalleled opportunity to investigate the neural, behavioral, and genetic underpinnings of child and adolescent development at scale (Marek et al., 2022; Wu et al., 2025; Garavan et al., 2018). Its multimodal neuroimaging assessments include structural MRI (sMRI), diffusion-weighted imaging (dMRI), and resting-state and task-based fMRI, along with extensive behavioral, cognitive, and environmental measures (Tomasi and Volkow, 2024).

A growing body of ABCD research has established links between brain morphology, functional connectivity, and behavioral or cognitive phenotypes such as psychopathology and general cognitive ability (Marek et al., 2022; Van Rooij et al., 2018). For instance, cortical thickness (CT) and functional connectivity (FC) have been linked to psychopathology (Child Behavior Checklist, or CBCL) and cognitive ability (NIH Toolbox total score) when modeled individually using support vector regression or canonical correlation analysis (CCA), but replicable associations typically require thousands of participants. Extending beyond single-modality analyses, recent multivariate frameworks integrating structural, diffusion, and functional MRI phenotypes have revealed shared genetic influences across imaging modalities (Elliott et al., 2018; Groves et al., 2011).

Other ABCD investigations have reported associations between higher body mass index (BMI) and reduced prefrontal cortical thickness (Laurent et al., 2020), altered cortical developmental trajectories in autism spectrum disorder (ASD) (Khundrakpam et al., 2017), and handedness-related lateralization of functional connectivity (Tomasi and Volkow, 2024). Collectively, these findings demonstrate the richness of the ABCD dataset for studying multivariate neural-behavioral relationships.

Despite these advances, most ABCD analyses remain modality-specific, emphasizing specific regions or univariate associations that require aggressive multiple-comparison correction and offer limited insight into cross-modality effects. When multimodal analyses are attempted, they typically rely on concatenation or dimension-reduction techniques (e.g., PCA, CCA), which merge feature spaces but do not decompose how distinct modalities uniquely or jointly contribute to group or behavioral differences (Joo et al., 2025). Furthermore, these approaches lack a unified inferential framework that can incorporate both continuous and categorical covariates or predictors, yield interpretable regression coefficients, and scale efficiently to large cohorts.

To address these challenges, we applied the proposed UGEE framework for distance-based inference to the ABCD dataset (release 5.0) (Garavan et al., 2018). Following the preprocessing pipeline of Kang et al. (2024), we extracted three imaging modalities: cortical thickness (CT) and surface area (SA) from structural MRI, and functional connectivity (FC) from resting-state fMRI involving subcortical regions. Each modality was represented by a pairwise distance matrix, constructed from Euclidean distance for CT and SA and distance of correlation vectors for FC, all centered and scaled for comparability.

After excluding diagnosis groups with less than 10 participants (i.e., intellectual disability or schizophrenia) and after strict image quality control (Kang et al., 2024), we included 8,543 individuals (1,341 disorders and 7,202 cognitively normal controls) for the analyses. The primary goal was to determine whether individuals with psychiatric disorders differed from cognitively normal (CN) individuals in any of the three imaging modalities. We defined the main predictor as a binary disorder vs. CN indicator and adjusted for BMI, handedness, cognitive composite score, and CBCL total problems score. Continuous covariates were centered and scaled prior to analysis. We also demonstrated the distance regression on more detailed disorder conditions in the Supplement as the secondary analyses. To eliminate site effect, we further conducted sensitivity analyses by regressing out site differences within each imaging modality and then repeating this distance regression.

### 4.2. ABCD Results

We first visualized modality-specific variation using principal coordinate analysis (PCoA; Figure 4(a)). Across CT, SA, and FC, the two groups showed minimal separation in the cluster centroid; however, the group with psychiatric conditions showed a greater spread, particularly in CT and FC.

**FIG 4.**
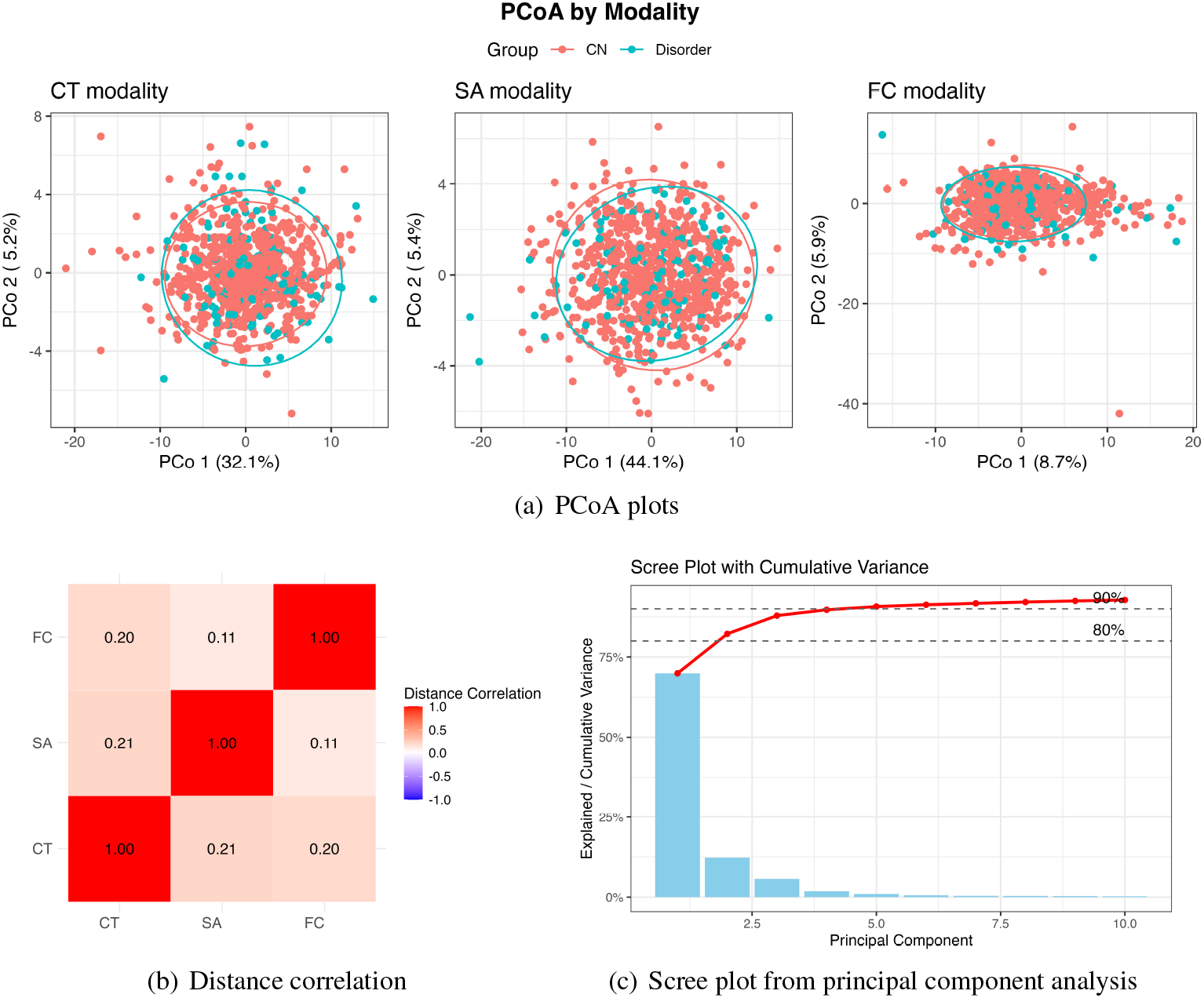
Exploratory analysis based on a 10% balanced subsample. (A) PCoA plots for cortical thickness (CT), surface area (SA), and functional connectivity (FC) modalities. CN (red) and disorder (blue) groups show substantial overlap. (B) Distance correlations among CT, SA, and FC. (C) Scree plot from principal component analysis on the averaged modality.

To quantify relationships among modalities, we computed pairwise distance correlations (Székely, Rizzo and Bakirov, 2007; Székely and Rizzo, 2009) as shown in Figure 4(b). The positive but moderate correlations across three modalities (0.11-0.21) implied that each may capture related yet distinct aspects of brain organization. Finally, principal component analysis (PCA) on the aggregated distance matrices (Figure 4(c)) showed that the first five components explained over 90% of the total variance; therefore, the top five PCs (*L* = 5) were used for the SiMMR-PC analysis.

We first compared SiMMR with three variants of the proposed distance regression: UGEE-Uni (a single aggregated distance), UGEE-Multi (independent working correlation), and UGEE-Multi (correlated). The inter-modality correlations were empirically estimated as in Figure 4(b). For each approach, we specified three models:

(1) A main-effect model;
(2) An interaction model including group-by-CBCL scores interaction;
(3) An interaction model including group-by-modality interaction (UGEE-Multi only). Using a 10% group-stratified subsample (*n* = 854; Table 2), we first assessed computational scalability across methods. This subsample size ensures that all methods, including SiMMR, could be executed within reasonable time and resource constraints. Under this setting, UGEE methods exhibited substantially lower computational cost while maintaining stable inference, with runtimes approximately 100-fold shorter than those of SiMMR (e.g., 5.68 seconds for UGEE-Multi vs. 10.6 minutes for SiMMR). Given these substantial differences in computational efficiency, we focused on UGEE-Uni and UGEE-Multi in subsequent analyses.

**Table 2.** Two omnibus p-values (omnibus and without-modality omnibus) and computation times (unit = sec) using a 10% subsample of the data (n = 854). The omnibus test (UGEE-Multi only) evaluates all main and interaction effects jointly; the without-modality omnibus test assesses all main and interaction effects other than the modality effects, which are directly comparable between UGEE-Uni and UGEE-Multi.

|  |  | SiMMR-D | SiMMR-PC | UGEE-Uni | UGEE-Multi (indep) | UGEE-Multi (corr) |
| --- | --- | --- | --- | --- | --- | --- |
| Main-effect model | p (omnibus) | — | — | — | 0.000 | 0.000 |
|  | p (omnibus_nomodality) | 0.009 | 0.009 | 0.034 | 0.034 | 0.048 |
|  | time (in second) | 636 | 636 | 0.33 | 5.68 | 5.47 |
| Interaction model<br>(Group × CBCL) | p (omnibus) | — | — | — | 0.000 | 0.000 |
|  | p (omnibus_nomodality) | 0.009 | 0.009 | 0.050 | 0.050 | 0.073 |
|  | time (in second) | 549 | 549 | 0.56 | 8.60 | 8.50 |

We then repeated the UGEE analyses on all available data (*n*=8,543). UGEE-Uni and UGEE-Multi yielded largely consistent conclusions across the specified models (Tables 3, 4 and Table S4 in the Supplement). Both modality contrasts were significant, indicating that beyond CT, FC and SA each explain additional variation in the between-subject dissimilarities, with FC showing a much larger effect size (Table 3 and 4). Interestingly, UGEE-Multi revealed modality differences in opposite directions: the SA-CT contrast was negative, whereas the FC-CT effect was positive. Since the distance regression parameterizes modality effects relative to CT (reference modality), a negative SA-CT coefficient indicates that between-subject dissimilarities in SA are smaller than those in CT. The strongly positive FC-CT effect implies larger between-subject dissimilarities in FC relative to CT, indicating that FC contributes more substantially to explaining multimodal variability across individuals.

**Table 3.** Group × CBCL interaction model results (coefficients, standardized effect size with p-values) with all available data (n=8,543). CT = cortical thickness; SA = surface area; FC = functional connectivity.

| Variable | UGEE-Uni |  |  | UGEE-Multi (indep) |  |  | UGEE-Multi (corr) |  |  |
| --- | --- | --- | --- | --- | --- | --- | --- | --- | --- |
|  | Coef. | Effect Size | p | Coef. | Effect Size | p | Coef. | Effect Size | p |
| ModalitySA_CT | – | – | – | -0.0099 | -6.321 | 0.016 | -0.0099 | -6.321 | 0.016 |
| ModalityFC_CT | – | – | – | 0.6388 | 299.567 | 0.000 | 0.6388 | 299.567 | 0.000 |
| Scale | 0.0526 | 6.822 | 0.000 | 0.0526 | 6.822 | 0.000 | 0.0546 | 6.262 | 0.000 |
| Location | 0.0324 | 12.328 | 0.000 | 0.0324 | 12.328 | 0.000 | 0.0336 | 11.418 | 0.000 |
| BMI | 0.0006 | 7.402 | 0.533 | 0.0006 | 7.402 | 0.533 | 0.0003 | 3.993 | 0.720 |
| Handedness | 0.0068 | 14.116 | 0.003 | 0.0068 | 14.116 | 0.003 | 0.0070 | 12.963 | 0.004 |
| Cognitive ability | 0.0012 | 7.294 | 0.378 | 0.0012 | 7.294 | 0.378 | 0.0011 | 6.489 | 0.416 |
| Psychopathology | 0.0083 | 23.311 | 0.000 | 0.0083 | 23.311 | 0.000 | 0.0088 | 21.543 | 0.000 |
| Scale × Psychopathology | -0.0056 | -2.760 | 0.230 | -0.0056 | -2.760 | 0.230 | -0.0058 | -2.482 | 0.250 |
| Location × Psychopathology | -0.0079 | -9.457 | 0.008 | -0.0079 | -9.457 | 0.008 | -0.0082 | -8.551 | 0.011 |
| Omnibus | – | – | – | – | – | 0.000 | – | – | 0.000 |
| omnibus_nomodality | – | – | 0.000 | – | – | 0.000 | – | – | 0.000 |

**Table 4.** Group × modality interaction model results (coefficients, standardized effect size with p-values) with all available data (n=8,543). CT = cortical thickness; SA = surface area; FC = functional connectivity.

| Variable | UGEE-Multi (indep) |  |  | UGEE-Multi (corr) |  |  |
| --- | --- | --- | --- | --- | --- | --- |
|  | Coef. | Effect Size | p | Coef. | Effect Size | p |
| ModalitySA_CT | -0.0077 | -4.242 | 0.083 | -0.0077 | -4.242 | 0.083 |
| ModalityFC_CT | 0.6336 | 268.068 | 0.000 | 0.6336 | 268.068 | 0.000 |
| Scale | 0.0350 | 5.084 | 0.000 | 0.0350 | 5.083 | 0.000 |
| Location | 0.0162 | 8.578 | 0.000 | 0.0161 | 8.508 | 0.000 |
| BMI | 0.0005 | 6.971 | 0.557 | 0.0003 | 3.583 | 0.747 |
| Handedness | 0.0069 | 14.139 | 0.003 | 0.0069 | 12.981 | 0.004 |
| Cognitive ability | 0.0011 | 7.255 | 0.380 | 0.0010 | 6.432 | 0.420 |
| Psychopathology | 0.0054 | 22.725 | 0.001 | 0.0057 | 21.363 | 0.001 |
| ModalitySA_CT × Scale | -0.0137 | -1.053 | 0.249 | -0.0137 | -1.053 | 0.249 |
| ModalitySA_CT × Location | -0.0069 | -2.071 | 0.248 | -0.0069 | -2.071 | 0.248 |
| ModalityFC_CT × Scale | 0.0307 | 1.539 | 0.037 | 0.0307 | 1.539 | 0.037 |
| ModalityFC_CT × Location | 0.0164 | 3.012 | 0.033 | 0.0164 | 3.012 | 0.033 |

Significant scale and location effects were also consistently detected across all UGEE variants, indicating that both group-level variability (scale) and mean differences (location) contribute substantially to the observed multimodal dissimilarities, which are consistent with the PCoA plot in Figure 4(a). We also noticed significant associations in location by CBCL total scores interaction in Table 3, between imaging features and behavioral measures, particularly for the group (scale and location effects) in the FC modality in Table 4, and replicated significant effects for handedness and CBCL total scores. UGEE-Multi uniquely detected subtle but significant cross-modality effects among CT, SA, and FC, indicating that structural and functional features contribute complementary information to diagnostic group differences.

We compared computational time in Table 5, where the expected trade-off between model complexity and runtime is evident. Secondary analyses for all four detailed disorder groups (Tables S5, S6) are included in the Supplement. Sensitivity analyses removing the site effects (Table S8, S9) are largely consistent with the primary results.

**Table 5.** Computation time with all available data (n=8,543; unit = sec).

|  | UGEE-Uni | UGEE-Multi (indep) | UGEE-Multi (corr) |
| --- | --- | --- | --- |
| Main-effect model | 32.27 | 611.70 | 571.10 |
| Interaction model (Group × CBCL) | 38.39 | 612.50 | 615.90 |
| Interaction model (Group × modality) | – | 779.54 | 812.03 |

These findings highlight UGEE’s ability to provide an efficient inferential framework for large-scale multimodal neuroimaging analysis. Empirically, the results are consistent with previous ABCD findings linking atypical cortical trajectories to ASD (Wu et al., 2025), handedness to the lateralization of multimodal brain patterns (Tomasi and Volkow, 2024), and CBCL to variability in both brain structure and functional connectivity (Marek et al., 2022). Methodologically, UGEE achieves substantial computational efficiency while yielding interpretable regression coefficients that support hypothesis-driven inference on modality-specific and cross-modality effects, which are not available in conventional distance-based approaches such as SiMMR or MDMR.

## 5. Discussion

The central goal of this study was to develop and apply a semiparametric, U-statistics-based Generalized Estimating Equation (UGEE) framework for distance-based inference that unifies univariate and multivariate distance regressions to enable interpretable, scalable, and statistically rigorous multimodal analyses. Using data from the ABCD study, we empirically demonstrated that UGEE achieves both computational efficiency and inferential robustness over existing nonparametric approaches.

Firstly, UGEE yields consistent and interpretable inference across both univariate and multivariate parameterizations, producing stable parameter estimates and robust *p*-values in large-scale analyses. This reproducibility supports recent calls in the neuroimaging literature for jointly analyzing phenotypes from multiple imaging sources and moving beyond single-modality frameworks (Marek et al., 2022; Tissink et al., 2024). In addition, UGEE achieves substantial computational efficiency without relying on permutation-based inference, completing analyses roughly 100 times faster than existing distance-based methods while maintaining asymptotic efficiency by virtue of the efficient influence function (EIF). This improvement ensures scalability to the increasingly high-dimensional data typical of large neuroimaging consortia.

Importantly, UGEE-Multi explicitly models modality-specific effects, which substantially improves statistical power and reduces required sample sizes, particularly when intermodality correlations are high. For example, under the simulation scenario with additive modality and scale (dispersion) effect, achieving 80% power to detect the target omnibus standardized effect size of 3 required *n* = 211 for aggregated models (UGEE-Uni) vs. *n* = 131 for modality-specific models (UGEE-Multi), based on the assumed linear relationship between standardized effect size and 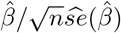. Similarly, when the true effect involved the modality and location (mean) differences, attaining 80% power to detect a target omnibus standardized effect size of 5 required *n* = 132 for UGEE-Uni but only *n* = 89 for UGEE-Multi, a one-third reduction. These findings highlight the efficiency gains achieved by explicitly modeling modality effects rather than collapsing them. Furthermore, UGEE-Multi captures cross-modality interactions, such as group-by-modality and covariate-by-modality effects, which remain inaccessible to aggregated or ANOVA-type frameworks.

Among existing multimodality integration methods, UGEE is conceptually related to Similarity Network Fusion (SNF) (Wang et al., 2014), which merges sample-level (dis)similarity matrices (or networks) across modalities for unsupervised clustering. Distinct from their goal of identifying latent subtypes, UGEE focuses on a hypothesis-driven and supervised inferential framework and enables formal testing of group, behavioral, or even genetic effects across multiple distance modalities. Other alternatives for multivariate analyses, such as Canonical Correlation Analysis (CCA) and its sparse variants (Gao, Ma and Zhou, 2017), can identify maximally correlated components across modalities but rely on linear and Gaussian assumptions, producing loadings rather than interpretable effect estimates. In contrast, UGEE directly specifies a semiparametric regression on pairwise distances, thereby capturing nonlinear dependencies while enabling asymptotic inference and interpretable effect size estimation.

Beyond neuroimaging, UGEE holds promise for uncovering brain-behavior pathways during preadolescence. Recent imaging-genetics studies using sparse CCA (Joo et al., 2025) have revealed bidirectional associations between polygenic scores (PGS) and neural organization: positive for cognitive-related PGS and structural MRI features, and negative for health-risk traits such as CBCL total problems across modalities, but these models are limited in capturing nonlinear or hierarchical dependencies. By directly modeling pairwise dissimilarities without assuming linearity or Gaussianity, UGEE may potentially offer a flexible, distribution-free alternative to sparse CCA for testing how polygenic signals propagate through multimodal brain structure and connectivity to shape behavioral phenotypes.

Several limitations warrant acknowledgment. First, although UGEE efficiently models inter-modality dependence, its performance may vary with the choice of distance metric and preprocessing quality, particularly for modalities with different signal-to-noise ratios. This sensitivity is inherent to distance-based approaches and could be mitigated through adaptive or domain-knowledge-guided metrics that better capture data-specific geometry (Reiss et al., 2010). Further, the current analysis focused on cross-sectional ABCD data; extending UGEE to longitudinal designs would enable modeling of within-subject developmental trajectories and temporal dependencies, yet the predominant missing data needs to be addressed explicitly in the semiparametric inference (Tsiatis, 2006).

Taken together, UGEE establishes a general inferential framework for multimodal data integration that combines statistical efficiency, interpretability, and scalability. By bridging distance-based ANOVA and regression paradigms, UGEE provides a unified statistical foundation for reproducible, hypothesis-driven multimodal inference, positioning it as a scalable tool for multimodal neuroimaging analyses. Codes of all analyses in this paper are available on GitHub: https://github.com/XinyuIvy/UGEE.git.

## Acknowledgments

The ABCD data used in this paper are from the NIMH Data Archive (https://doi.org/10.15154/1503209) and the ABCD BIDS Community Collection (ABCC; https://collection3165.readthedocs.io). The authors would like to thank the anonymous referees, an Associate Editor, and the Editor for their constructive comments that improved the quality of this paper.

## SUPPLEMENTARY MATERIAL

### Supplement

#### S5.1. Notations

Please see all notations in Table S1.

**Table S1.**
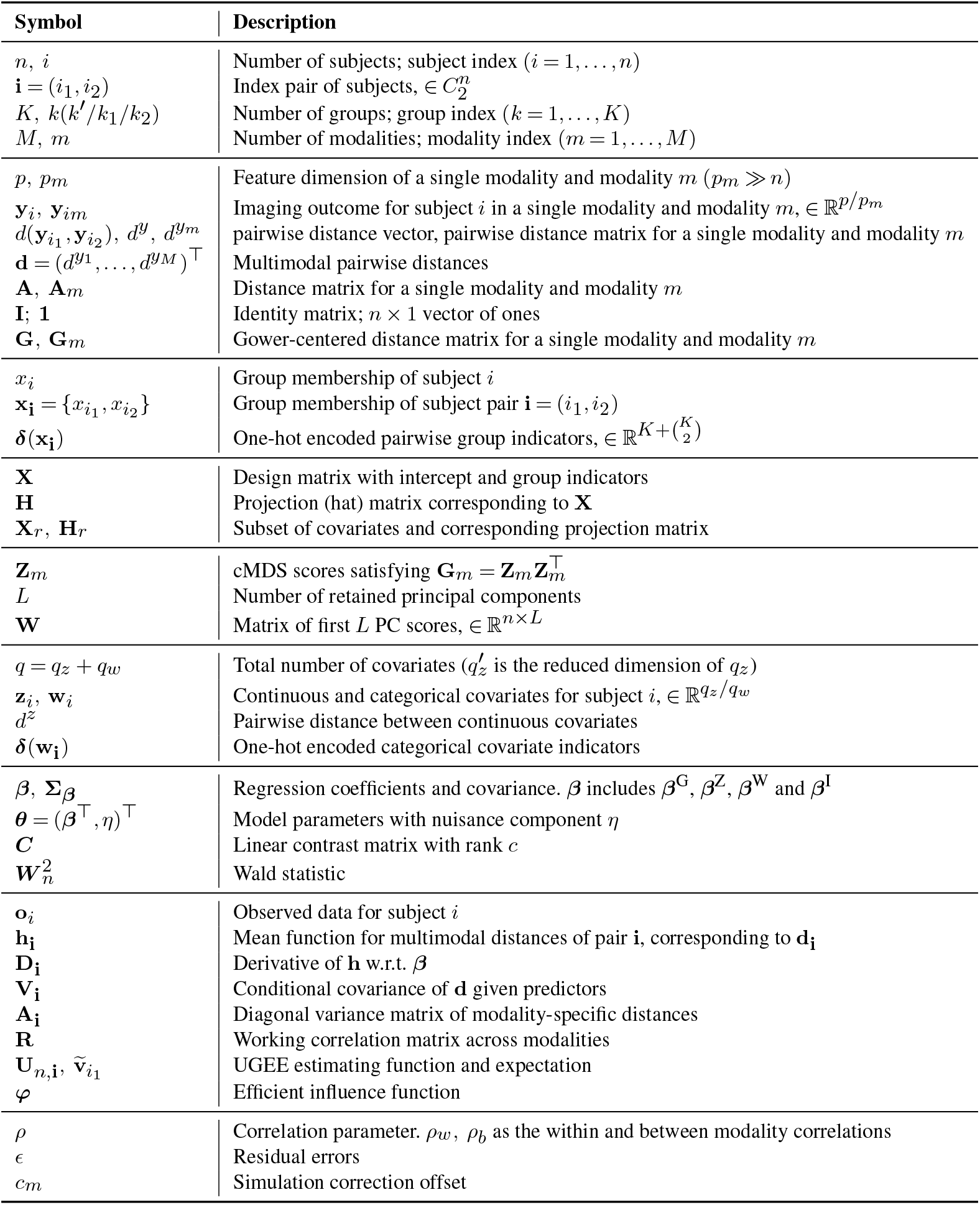
Notation summary.

| Symbol | Description |
| --- | --- |
| $n, i$<br>$\mathbf{i} = (i_1, i_2)$<br>$K, k(k'/k_1/k_2)$<br>$M, m$ | Number of subjects; subject index ($i = 1, \dots, n$)<br>Index pair of subjects, $\in C_2^n$<br>Number of groups; group index ($k = 1, \dots, K$)<br>Number of modalities; modality index ($m = 1, \dots, M$) |
| $p, p_m$<br>$\mathbf{y}_i, \mathbf{y}_{im}$<br>$d(\mathbf{y}_{i_1}, \mathbf{y}_{i_2}), d^y, d^{y_m}$<br>$\mathbf{d} = (d^{y_1}, \dots, d^{y_M})^\top$<br>$\mathbf{A}, \mathbf{A}_m$<br>$\mathbf{I}; \mathbf{1}$<br>$\mathbf{G}, \mathbf{G}_m$ | Feature dimension of a single modality and modality $m$ ($p_m \gg n$)<br>Imaging outcome for subject $i$ in a single modality and modality $m$, $\in \mathbb{R}^{p/p_m}$<br>pairwise distance vector, pairwise distance matrix for a single modality and modality $m$<br>Multimodal pairwise distances<br>Distance matrix for a single modality and modality $m$<br>Identity matrix; $n \times 1$ vector of ones<br>Gower-centered distance matrix for a single modality and modality $m$ |
| $x_i$<br>$\mathbf{x}_i = \{x_{i_1}, x_{i_2}\}$<br>$\delta(\mathbf{x}_i)$ | Group membership of subject $i$<br>Group membership of subject pair $\mathbf{i} = (i_1, i_2)$<br>One-hot encoded pairwise group indicators, $\in \mathbb{R}^{K + \binom{K}{2}}$ |
| $\mathbf{X}$<br>$\mathbf{H}$<br>$\mathbf{X}_r, \mathbf{H}_r$ | Design matrix with intercept and group indicators<br>Projection (hat) matrix corresponding to $\mathbf{X}$<br>Subset of covariates and corresponding projection matrix |
| $\mathbf{Z}_m$<br>$L$<br>$\mathbf{W}$ | cMDS scores satisfying $\mathbf{G}_m = \mathbf{Z}_m \mathbf{Z}_m^\top$<br>Number of retained principal components<br>Matrix of first $L$ PC scores, $\in \mathbb{R}^{n \times L}$ |
| $q = q_z + q_w$<br>$\mathbf{z}_i, \mathbf{w}_i$<br>$d^z$<br>$\delta(\mathbf{w}_i)$ | Total number of covariates ($q'_z$ is the reduced dimension of $q_z$)<br>Continuous and categorical covariates for subject $i$, $\in \mathbb{R}^{q_z/q_w}$<br>Pairwise distance between continuous covariates<br>One-hot encoded categorical covariate indicators |
| $\beta, \Sigma_\beta$<br>$\theta = (\beta^\top, \eta)^\top$<br>$\mathbf{C}$<br>$\mathbf{W}_n^2$ | Regression coefficients and covariance. $\beta$ includes $\beta^G, \beta^Z, \beta^W$ and $\beta^I$<br>Model parameters with nuisance component $\eta$<br>Linear contrast matrix with rank $c$<br>Wald statistic |
| $\mathbf{o}_i$<br>$\mathbf{h}_i$<br>$\mathbf{D}_i$<br>$\mathbf{V}_i$<br>$\mathbf{A}_i$<br>$\mathbf{R}$<br>$\mathbf{U}_{n,i}, \tilde{\mathbf{v}}_{i_1}$<br>$\varphi$ | Observed data for subject $i$<br>Mean function for multimodal distances of pair $\mathbf{i}$, corresponding to $\mathbf{d}_i$<br>Derivative of $\mathbf{h}$ w.r.t. $\beta$<br>Conditional covariance of $\mathbf{d}$ given predictors<br>Diagonal variance matrix of modality-specific distances<br>Working correlation matrix across modalities<br>UGEE estimating function and expectation<br>Efficient influence function |
| $\rho$<br>$\epsilon$<br>$c_m$ | Correlation parameter. $\rho_w, \rho_b$ as the within and between modality correlations<br>Residual errors<br>Simulation correction offset |

#### S5.2. Proof of Theorem 1

A *regular* and *asymptotically linear* (RAL) between-subject 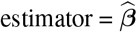 for distance-based regression satisfies

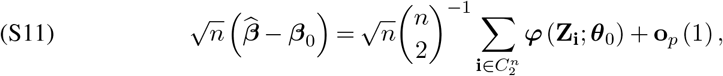

where 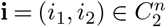 indexes all distinct subject pairs. The Efficient UGEE is defined in the Main paper satisfies (S11) by verifying that the influence function (IF) satisfies

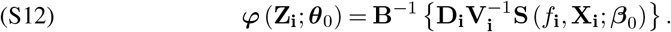

It is important to note that the pairwise influence functions ***φ*** (**Z**_**i**_; ***θ***_0_) in (S11) are *not* mutually independent. For instance, the pairwise vectors ***φ*** (**Z**_**i**_; ***θ***_0_) formed by (**Z**_1_, **Z**_2_) and (**Z**_1_, **Z**_3_) share the common subject **Z**_1_, thereby inducing interlocking correlations across them.

Although the summation in (S11) involves 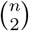 distinct pairs, the resulting estimator 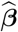 remains 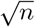 -consistent as we show below. Define a linear, many-to-one mapping by

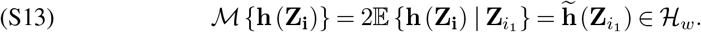

For any **h** (**Z**_**i**_), the mapping ℳ (·) projects this pairwise function onto the within-subject space via conditional expectation, creating an image 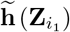 that depends only on one subject and thus forms an independent sequence across subjects.

Now, applying ℳ (·) in (S13) to the influence functions ***φ*** (**Z**_**i**_; ***θ***_0_)’s in (S11) breaks the dependencies among pairs. Along with the theory of U-statistics (Liu et al., 2024), this yields:

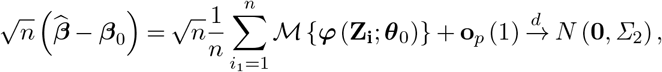

where **o**_*p*_ (1) denotes the stochastic version of **o** (1), and the asymptotic variance-covariance matrix for 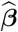 is:

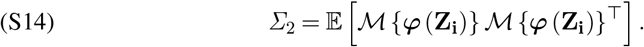

This establishes the 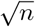 -consistency and asymptotic normality (AN) of any RAL estimator satisfying (S11).

Further, by a series of results based on Hilbert-space-based efficiency theory in Liu et al. (2022b), (S12) is shown to be the unique *efficient influence function* (EIF) whose asymptotic variance *Σ*_2_ achieves the semiparametric efficiency bound within the model class under the same moment restriction, which can be calculated as

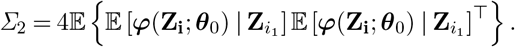

In practice, consistent estimators of *Σ*_2_ can be obtained by substituting consistent estimators of ***β*** and moment estimators of the respective quantities.

#### S5.3. Positive-definiteness of the block correlation matrix

Consider *M* modalities, each with *p* features, and a block compound symmetry correlation matrix **R**(*ρ*) with within-modality correlation *ρ*_*w*_ (off-diagonal elements within each block) and between-modality correlation *ρ*_*b*_ (elements across blocks), with unit diagonals. The eigenvalues of **R**(*ρ*) are

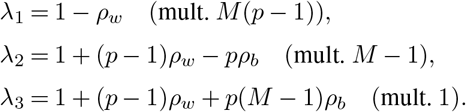

Hence, **R**(*ρ*) is positive definite if and only if

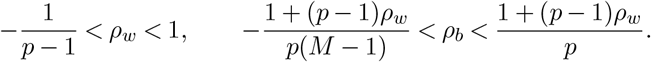

These conditions are both necessary and sufficient and do not require *ρ*_*b*_ ≤ *ρ*_*w*_; the admissible range of *ρ*_*b*_ depends on (*p, M, ρ*_*w*_).

#### S5.4. Bias correction c_*m*_ of Euclidean distances inside an exponential link

Under independent features within the *m* th modality, we have 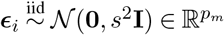 and the pairwise Euclidean distances

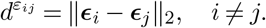

Then ***ϵ***_*i*_ − ***ϵ***_*j*_ ∼ *N* (**0**, 2*s*^2^**I**) and

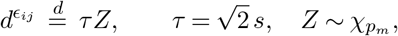

where 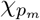 denotes the chi distribution with *p*_*m*_ degrees of freedom. Placing 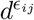 inside an exponential link inflates the mean, and we seek a constant *c* such that

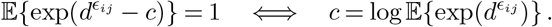

**Case** *p*_*m*_ = 1 **(scalar residuals)**. In this case 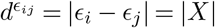 with *X* ∼ 𝒩 (0, *τ* ^2^) (a folded normal variable). A standard calculation gives

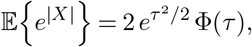

where Φ(·) is the standard normal cumulative distribution function. Substituting 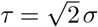 yields

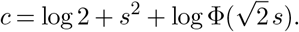

**Case** *p*_*m*_ ≥ 1 **(general Euclidean distance)**. Let *R* = *τ Z* with 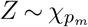. Then

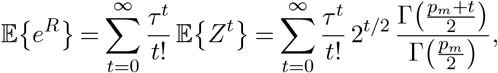

using the known moments of the chi distribution, 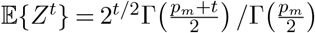. Therefore,

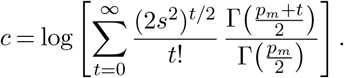

In the simulation, *t* is set from 0 to 100. Subtracting this constant from the exponent ensures that 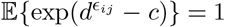, thus removing the artificial mean inflation caused by the distance-based residual term. For numerical implementation, the series converges rapidly for moderate *s*, and an additional normalization by the empirical mean of off-diagonal entries can be applied to further stabilize mean(*G*) ≈ 1.

#### S5.5. Additional simulation results

Tables S2 and S3 show the bias and standardized effect size for UGEE-Multi across different scenarios and sample sizes under independent and correlated features.

#### S5.6. Additional application results

**Table S2.**
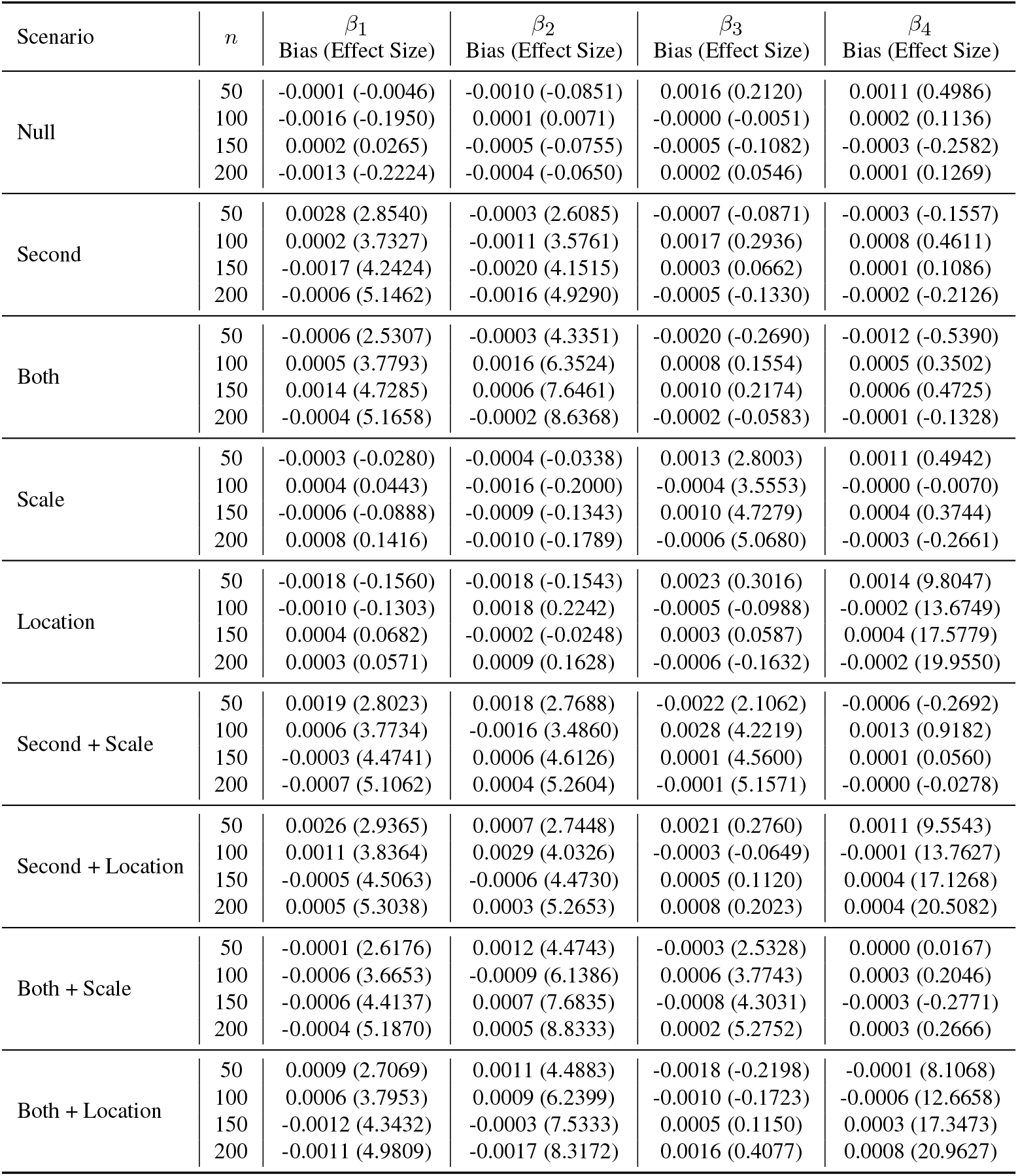
Bias and standardized effect size for UGEE-Multi across different scenarios and sample sizes under independent features (ρ_w_ = 0). UGEE-Uni shares the same bias as UGEE-Multi.

| Scenario | $n$ | $\beta_1$<br>Bias (Effect Size) | $\beta_2$<br>Bias (Effect Size) | $\beta_3$<br>Bias (Effect Size) | $\beta_4$<br>Bias (Effect Size) |
| --- | --- | --- | --- | --- | --- |
| Null | 50 | -0.0001 (-0.0046) | -0.0010 (-0.0851) | 0.0016 (0.2120) | 0.0011 (0.4986) |
|  | 100 | -0.0016 (-0.1950) | 0.0001 (0.0071) | -0.0000 (-0.0051) | 0.0002 (0.1136) |
|  | 150 | 0.0002 (0.0265) | -0.0005 (-0.0755) | -0.0005 (-0.1082) | -0.0003 (-0.2582) |
|  | 200 | -0.0013 (-0.2224) | -0.0004 (-0.0650) | 0.0002 (0.0546) | 0.0001 (0.1269) |
| Second | 50 | 0.0028 (2.8540) | -0.0003 (2.6085) | -0.0007 (-0.0871) | -0.0003 (-0.1557) |
|  | 100 | 0.0002 (3.7327) | -0.0011 (3.5761) | 0.0017 (0.2936) | 0.0008 (0.4611) |
|  | 150 | -0.0017 (4.2424) | -0.0020 (4.1515) | 0.0003 (0.0662) | 0.0001 (0.1086) |
|  | 200 | -0.0006 (5.1462) | -0.0016 (4.9290) | -0.0005 (-0.1330) | -0.0002 (-0.2126) |
| Both | 50 | -0.0006 (2.5307) | -0.0003 (4.3351) | -0.0020 (-0.2690) | -0.0012 (-0.5390) |
|  | 100 | 0.0005 (3.7793) | 0.0016 (6.3524) | 0.0008 (0.1554) | 0.0005 (0.3502) |
|  | 150 | 0.0014 (4.7285) | 0.0006 (7.6461) | 0.0010 (0.2174) | 0.0006 (0.4725) |
|  | 200 | -0.0004 (5.1658) | -0.0002 (8.6368) | -0.0002 (-0.0583) | -0.0001 (-0.1328) |
| Scale | 50 | -0.0003 (-0.0280) | -0.0004 (-0.0338) | 0.0013 (2.8003) | 0.0011 (0.4942) |
|  | 100 | 0.0004 (0.0443) | -0.0016 (-0.2000) | -0.0004 (3.5553) | -0.0000 (-0.0070) |
|  | 150 | -0.0006 (-0.0888) | -0.0009 (-0.1343) | 0.0010 (4.7279) | 0.0004 (0.3744) |
|  | 200 | 0.0008 (0.1416) | -0.0010 (-0.1789) | -0.0006 (5.0680) | -0.0003 (-0.2661) |
| Location | 50 | -0.0018 (-0.1560) | -0.0018 (-0.1543) | 0.0023 (0.3016) | 0.0014 (9.8047) |
|  | 100 | -0.0010 (-0.1303) | 0.0018 (0.2242) | -0.0005 (-0.0988) | -0.0002 (13.6749) |
|  | 150 | 0.0004 (0.0682) | -0.0002 (-0.0248) | 0.0003 (0.0587) | 0.0004 (17.5779) |
|  | 200 | 0.0003 (0.0571) | 0.0009 (0.1628) | -0.0006 (-0.1632) | -0.0002 (19.9550) |
| Second + Scale | 50 | 0.0019 (2.8023) | 0.0018 (2.7688) | -0.0022 (2.1062) | -0.0006 (-0.2692) |
|  | 100 | 0.0006 (3.7734) | -0.0016 (3.4860) | 0.0028 (4.2219) | 0.0013 (0.9182) |
|  | 150 | -0.0003 (4.4741) | 0.0006 (4.6126) | 0.0001 (4.5600) | 0.0001 (0.0560) |
|  | 200 | -0.0007 (5.1062) | 0.0004 (5.2604) | -0.0001 (5.1571) | -0.0000 (-0.0278) |
| Second + Location | 50 | 0.0026 (2.9365) | 0.0007 (2.7448) | 0.0021 (0.2760) | 0.0011 (9.5543) |
|  | 100 | 0.0011 (3.8364) | 0.0029 (4.0326) | -0.0003 (-0.0649) | -0.0001 (13.7627) |
|  | 150 | -0.0005 (4.5063) | -0.0006 (4.4730) | 0.0005 (0.1120) | 0.0004 (17.1268) |
|  | 200 | 0.0005 (5.3038) | 0.0003 (5.2653) | 0.0008 (0.2023) | 0.0004 (20.5082) |
| Both + Scale | 50 | -0.0001 (2.6176) | 0.0012 (4.4743) | -0.0003 (2.5328) | 0.0000 (0.0167) |
|  | 100 | -0.0006 (3.6653) | -0.0009 (6.1386) | 0.0006 (3.7743) | 0.0003 (0.2046) |
|  | 150 | -0.0006 (4.4137) | 0.0007 (7.6835) | -0.0008 (4.3031) | -0.0003 (-0.2771) |
|  | 200 | -0.0004 (5.1870) | 0.0005 (8.8333) | 0.0002 (5.2752) | 0.0003 (0.2666) |
| Both + Location | 50 | 0.0009 (2.7069) | 0.0011 (4.4883) | -0.0018 (-0.2198) | -0.0001 (8.1068) |
|  | 100 | 0.0006 (3.7953) | 0.0009 (6.2399) | -0.0010 (-0.1723) | -0.0006 (12.6658) |
|  | 150 | -0.0012 (4.3432) | -0.0003 (7.5333) | 0.0005 (0.1150) | 0.0003 (17.3473) |
|  | 200 | -0.0011 (4.9809) | -0.0017 (8.3172) | 0.0016 (0.4077) | 0.0008 (20.9627) |

##### S5.6.1. Primary analysis

Table S4 is the result of the main effect model with two groups.

##### S5.6.2. Secondary analysis

Among 8,543 individuals, we have 136 with ASD, 1,052 with other developmental disorders, 153 with other psychiatric disorders, and 7,202 cognitively normal controls. Other developmental disorders covers conditions such as intellectual disability, language disorders, learning disorders, and motor coordination disorders. These are distinct from ASD but still impact developmental trajectories. Other psychiatric disorders includes diagnoses such as ADHD, anxiety disorders, depressive disorders, and disruptive behavior disorders. Figure S1 is the PCoA plot with four groups. Tables S5 and S6 are the results of the main effect model and model with group *×* CBCL interaction with four groups.

**Table S3.** Bias and standardized effect size for UGEE-Multi across different scenarios and sample sizes under correlated features (ρ_w_ = 0.8). UGEE-Uni shares the same bias as UGEE-Multi.

| Scenario | $n$ | $\beta_1$<br>Bias (Effect Size) | $\beta_2$<br>Bias (Effect Size) | $\beta_3$<br>Bias (Effect Size) | $\beta_4$<br>Bias (Effect Size) |
| --- | --- | --- | --- | --- | --- |
| Null | 50 | 0.0011 (0.1850) | -0.0007 (-0.1231) | 0.0008 (0.0380) | 0.0010 (0.1591) |
|  | 100 | -0.0005 (-0.1239) | 0.0012 (0.2984) | 0.0028 (0.1807) | 0.0014 (0.3437) |
|  | 150 | 0.0006 (0.1904) | 0.0016 (0.4777) | -0.0010 (-0.0803) | -0.0002 (-0.0534) |
|  | 200 | 0.0004 (0.1556) | 0.0008 (0.2806) | 0.0021 (0.1928) | 0.0013 (0.4505) |
| Second | 50 | -0.0023 (4.9154) | -0.0025 (4.7783) | 0.0026 (0.1145) | 0.0011 (0.1660) |
|  | 100 | -0.0013 (7.0780) | -0.0010 (7.0275) | -0.0026 (-0.1644) | -0.0009 (-0.2067) |
|  | 150 | 0.0007 (9.2185) | 0.0015 (9.5080) | -0.0001 (-0.0094) | 0.0001 (0.0348) |
|  | 200 | -0.0003 (10.2801) | -0.0000 (10.3429) | -0.0008 (-0.0746) | -0.0004 (-0.1369) |
| Both | 50 | -0.0013 (5.1646) | -0.0015 (8.5832) | -0.0016 (-0.0723) | -0.0005 (-0.0770) |
|  | 100 | 0.0006 (7.4959) | 0.0016 (12.6846) | 0.0036 (0.2286) | 0.0023 (0.5266) |
|  | 150 | -0.0018 (8.5062) | -0.0010 (14.5598) | -0.0021 (-0.1609) | -0.0007 (-0.2081) |
|  | 200 | -0.0007 (10.2253) | -0.0002 (17.2573) | 0.0010 (0.0914) | 0.0004 (0.1342) |
| Scale | 50 | -0.0017 (-0.2952) | -0.0016 (-0.2838) | -0.0008 (0.9070) | 0.0003 (0.0547) |
|  | 100 | 0.0001 (0.0172) | 0.0001 (0.0220) | -0.0018 (1.1512) | -0.0008 (-0.1868) |
|  | 150 | 0.0008 (0.2525) | 0.0001 (0.0216) | -0.0001 (1.5287) | 0.0001 (0.0308) |
|  | 200 | 0.0003 (0.1057) | -0.0005 (-0.1668) | 0.0011 (1.8985) | 0.0006 (0.2041) |
| Location | 50 | 0.0020 (0.3620) | 0.0022 (0.3888) | 0.0044 (0.2003) | 0.0027 (3.6135) |
|  | 100 | 0.0019 (0.4609) | 0.0008 (0.1913) | -0.0016 (-0.1011) | -0.0005 (4.6563) |
|  | 150 | -0.0001 (-0.0345) | -0.0004 (-0.1365) | -0.0022 (-0.1754) | -0.0011 (5.6825) |
|  | 200 | -0.0010 (-0.3385) | -0.0008 (-0.2670) | 0.0019 (0.1677) | 0.0011 (7.1584) |
| Second + Scale | 50 | 0.0021 (5.7226) | 0.0018 (5.6845) | 0.0014 (1.0051) | 0.0016 (0.2550) |
|  | 100 | 0.0009 (7.6189) | -0.0008 (7.2403) | -0.0050 (0.9528) | -0.0022 (-0.5228) |
|  | 150 | 0.0001 (9.0174) | 0.0003 (9.1483) | -0.0019 (1.4207) | -0.0008 (-0.2538) |
|  | 200 | 0.0005 (10.6977) | 0.0010 (10.8058) | -0.0002 (1.7831) | -0.0001 (-0.0371) |
| Second + Location | 50 | 0.0001 (5.2005) | -0.0007 (5.1661) | 0.0004 (0.0193) | 0.0002 (3.2151) |
|  | 100 | 0.0006 (7.5632) | -0.0002 (7.3034) | -0.0006 (-0.0395) | -0.0001 (4.7469) |
|  | 150 | -0.0004 (8.9586) | -0.0009 (8.8131) | 0.0004 (0.0302) | 0.0006 (6.0887) |
|  | 200 | -0.0003 (10.3081) | -0.0006 (10.2669) | -0.0010 (-0.0885) | -0.0004 (6.7361) |
| Both + Scale | 50 | -0.0003 (5.1605) | 0.0001 (8.6833) | -0.0019 (0.8237) | -0.0004 (-0.0551) |
|  | 100 | 0.0005 (7.3849) | 0.0004 (12.3749) | 0.0004 (1.2835) | 0.0007 (0.1511) |
|  | 150 | 0.0008 (9.2746) | 0.0009 (15.5127) | 0.0026 (1.7638) | 0.0016 (0.4624) |
|  | 200 | 0.0011 (10.6907) | 0.0005 (17.5949) | 0.0011 (1.8902) | 0.0008 (0.2786) |
| Both + Location | 50 | -0.0019 (4.9718) | -0.0003 (8.8562) | 0.0029 (0.1379) | 0.0024 (3.6413) |
|  | 100 | -0.0001 (7.3636) | 0.0013 (12.5467) | -0.0025 (-0.1581) | -0.0011 (4.4680) |
|  | 150 | -0.0009 (8.6734) | -0.0008 (14.8360) | -0.0001 (-0.0036) | 0.0003 (6.0093) |
|  | 200 | -0.0012 (10.0087) | -0.0007 (16.9496) | -0.0021 (-0.1931) | -0.0011 (6.5898) |

##### S5.6.3. Sensitivity analysis

Tables S8 and S9 are the results of the main effect model and model with group *×* CBCL interaction, after removing the site effects.

**Table S4.**
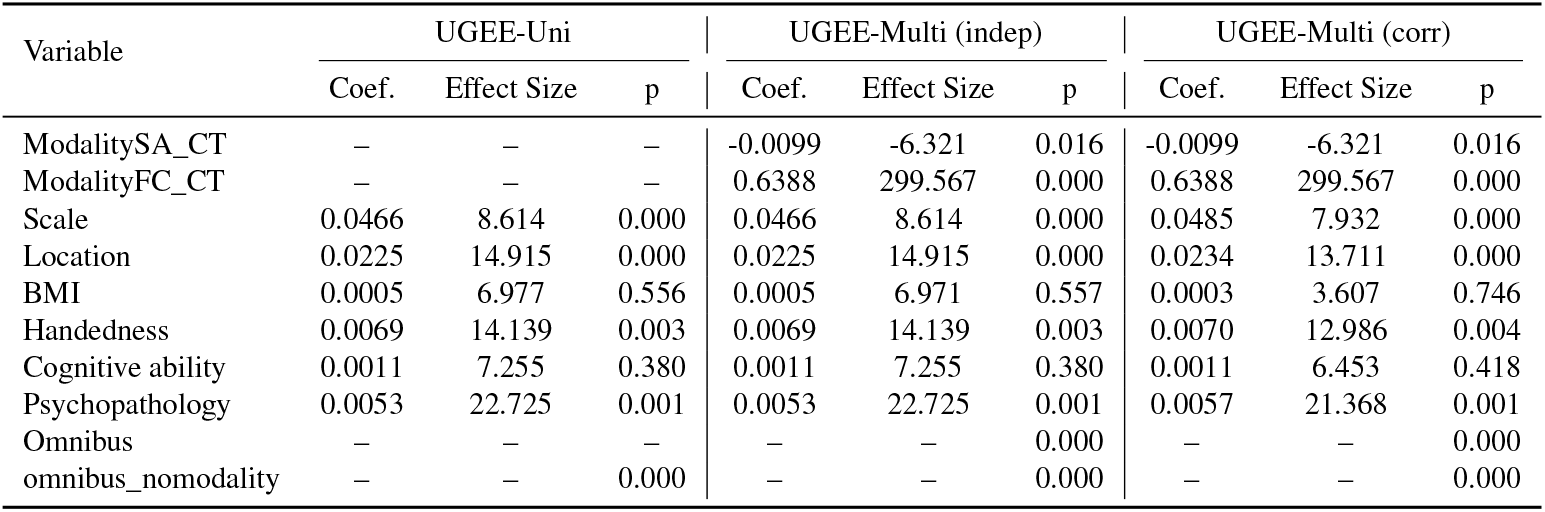
Main-effect model results (coefficients, standardized effect size with p-values) with all available data (n=8,543). CT = cortical thickness; SA = surface area; FC = functional connectivity.

**FIG S1.**
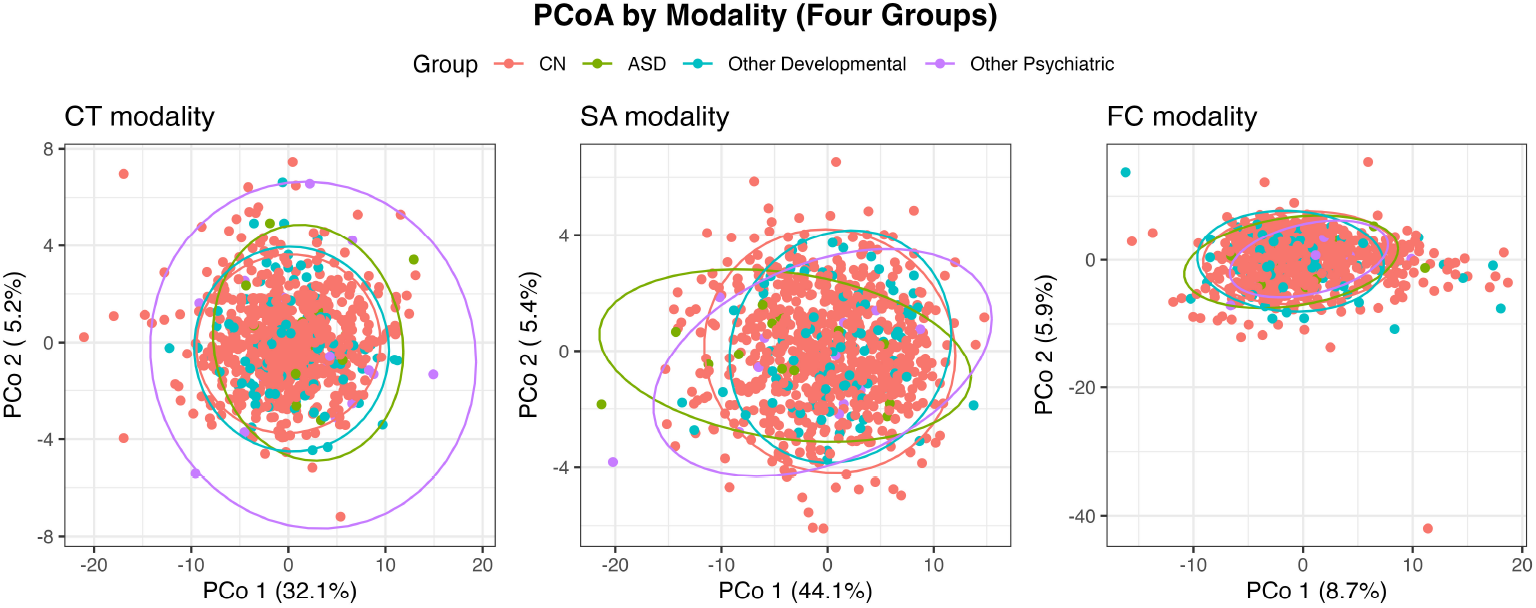
PCoA plots with four groups for cortical thickness (CT), surface area (SA), and functional connectivity (FC) modalities.

**Table S5.**
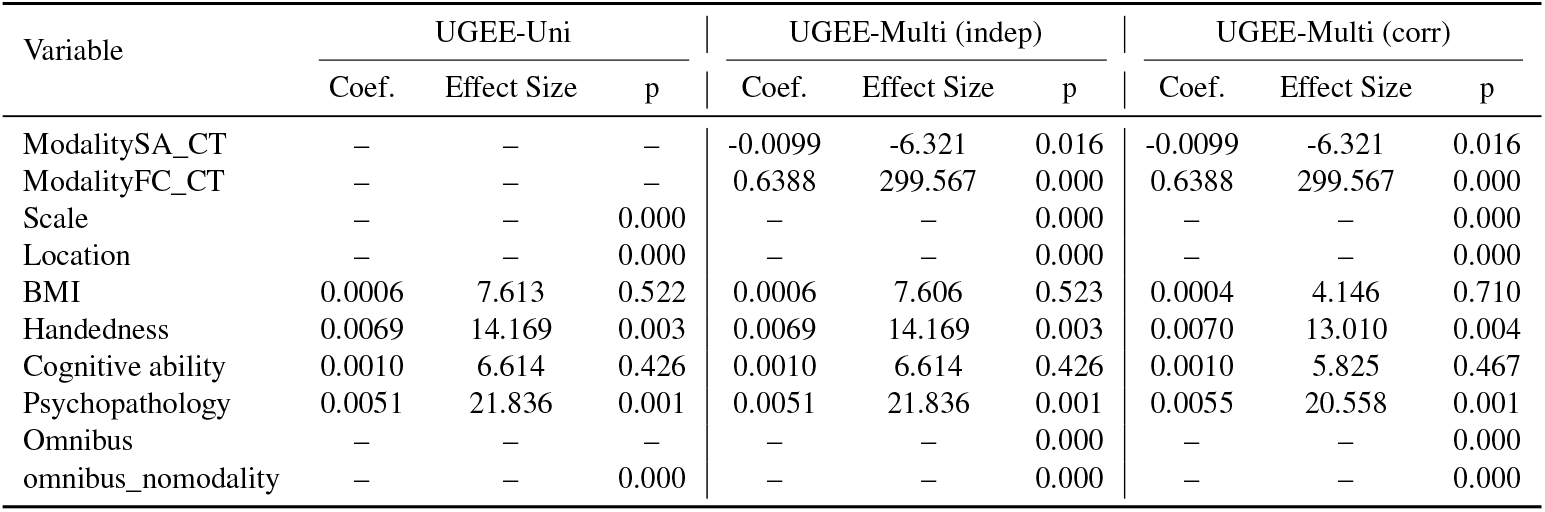
Main-effect model results with four groups (coefficients, standardized effect size with p-values) using all available data (n=8,543). CT = cortical thickness; SA = surface area; FC = functional connectivity.

**Table S6.** Group × CBCL interaction model results with four groups (coefficients, standardized effect size with p-values) using all available data (n=8,543).

| Variable | UGEE-Uni |  |  | UGEE-Multi (indep) |  |  | UGEE-Multi (corr) |  |  |
| --- | --- | --- | --- | --- | --- | --- | --- | --- | --- |
|  | Coef. | Effect Size | p | Coef. | Effect Size | p | Coef. | Effect Size | p |
| ModalitySA_CT | – | – | – | -0.0099 | -6.321 | 0.016 | -0.0099 | -6.321 | 0.016 |
| ModalityFC_CT | – | – | – | 0.6388 | 299.567 | 0.000 | 0.6388 | 299.567 | 0.000 |
| Scale | – | – | 0.000 | – | – | 0.000 | – | – | 0.000 |
| Location | – | – | 0.000 | – | – | 0.000 | – | – | 0.000 |
| BMI | 0.0006 | 8.085 | 0.497 | 0.0006 | 8.085 | 0.497 | 0.0004 | 4.578 | 0.682 |
| Handedness | 0.0069 | 14.157 | 0.003 | 0.0069 | 14.157 | 0.003 | 0.0070 | 12.995 | 0.004 |
| Cognitive ability | 0.0010 | 6.599 | 0.428 | 0.0010 | 6.599 | 0.428 | 0.0010 | 5.813 | 0.468 |
| Psychopathology | 0.0083 | 23.311 | 0.000 | 0.0083 | 23.311 | 0.000 | 0.0088 | 21.543 | 0.000 |
| Scale × Psychopathology | – | – | 0.278 | – | – | 0.278 | – | – | 0.326 |
| Location × Psychopathology | – | – | 0.150 | – | – | 0.150 | – | – | 0.183 |
| Omnibus | – | – | 0.000 | – | – | 0.000 | – | – | 0.000 |
| omnibus_nomodality | – | – | 0.000 | – | – | 0.000 | – | – | 0.000 |

**Table S7.** Computation time with all available data (unit = sec) with four groups.

|  | UGEE-Uni | UGEE-Multi (indep) | UGEE-Multi (corr) |
| --- | --- | --- | --- |
| Main-effect model | 49.49 | 707.40 | 700.70 |
| Interaction model (Group × CBCL) | 96.32 | 893.60 | 892.30 |

**Table S8.** Main-effect model results (coefficients, standardized effect size with p-values) with all available data (n=8,543). CT = cortical thickness; SA = surface area; FC = functional connectivity.

| Variable | UGEE-Uni |  |  | UGEE-Multi (indep) |  |  | UGEE-Multi (corr) |  |  |
| --- | --- | --- | --- | --- | --- | --- | --- | --- | --- |
|  | Coef. | Effect Size | p | Coef. | Effect Size | p | Coef. | Effect Size | p |
| ModalitySA_CT | – | – | – | 0.005 | 3.299 | 0.214 | 0.005 | 3.299 | 0.214 |
| ModalityFC_CT | – | – | – | 0.651 | 307.829 | 0.000 | 0.651 | 307.829 | 0.000 |
| Scale | 0.048 | 8.882 | 0.000 | 0.048 | 8.882 | 0.000 | 0.050 | 8.088 | 0.000 |
| Location | 0.023 | 15.265 | 0.000 | 0.023 | 15.265 | 0.000 | 0.024 | 13.897 | 0.000 |
| BMI | 0.001 | 7.154 | 0.552 | 0.001 | 7.147 | 0.553 | 0.000 | 3.293 | 0.769 |
| Handedness | 0.007 | 13.942 | 0.003 | 0.007 | 13.942 | 0.003 | 0.007 | 12.791 | 0.004 |
| Cognitive ability | 0.002 | 10.534 | 0.207 | 0.002 | 10.534 | 0.207 | 0.002 | 9.202 | 0.249 |
| Psychopathology | 0.006 | 24.055 | 0.000 | 0.006 | 24.055 | 0.000 | 0.006 | 22.230 | 0.000 |
| Omnibus | – | – | – | – | – | 0.000 | – | – | 0.000 |
| omnibus_nomodality | – | – | 0.000 | – | – | 0.000 | – | – | 0.000 |

**Table S9.**
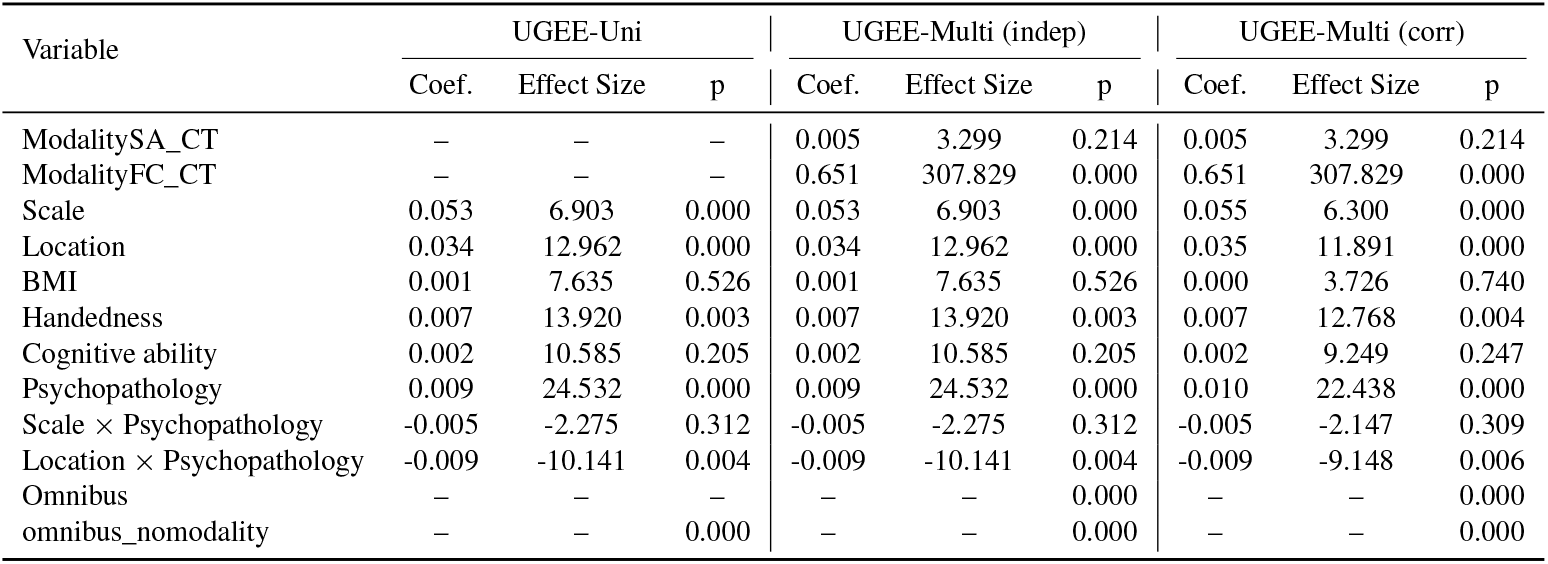
Group × CBCL interaction model results after removing site effects (coefficients, standardized effect size with p-values) with all available data (n=8,543). CT = cortical thickness; SA = surface area; FC = functional connectivity.

